# Mapping axon conduction delays *in vivo* from microstructural MRI

**DOI:** 10.1101/503763

**Authors:** Mark Drakesmith, Derek K Jones

**Affiliations:** Cardiff University Brain Research Imaging Centre, Cardiff University, Cardiff, UK; Neuroscience and Mental Health Research Institute, Cardiff University, Cardiff, UK; Mary McKillop Institute for Health Research, Faculty of Health Sciences, Australian Catholic University, Melbourne, Victoria 3065, Australia

## Abstract

The conduction velocity (CV) of action potentials along axons is a key neurophysiological property central to neural communication. The ability to estimate CV in humans *in vivo* from non-invasive MRI methods would therefore represent a significant advance in neuroscience. However, there are 2 major challenges that this paper aims to address: (1) much of the complexity of the neurophysiology of action potentials cannot be captured with currently available MRI techniques. Therefore, we seek to establish the variability in CV that *can* be captured when predicting CV purely from parameters that can be estimated from MRI (axon diameter and g-ratio); and (2) errors inherent in existing MRI-based biophysical models of tissue will propagate through to estimates of CV, the extent to which is currently unknown.

Issue (1) is investigated by performing a sensitivity analysis on a comprehensive model of axon electrophysiology and determining the relative sensitivity to various morphological and electrical parameters.

The investigations suggest that 89.2 % of the variance in CV is accounted for by variation in AD and g-ratio. The observed dependency of CV on AD and g-ratio is well characterised by a previously reported model by Rushton.

Issue (2) is investigated through simulation of diffusion and relaxometry MRI data for a range of axon morphologies, applying models of restricted diffusion and relaxation processes to derive estimates of axon volume fraction (AVF), AD and g-ratio and estimating CV from the derived parameters. The results show that errors in the AVF have the biggest detrimental impact on estimates of CV, particularly for sparse fibre populations (AVF*<* 0.3). CV estimates are most accurate (below 5% error) where AVF is above 0.3, g-ratio is between 0.6 and 0.85 and AD is below 10 µm. Fortunately, these parameter bounds are typically satisfied by most myelinated axons.

In conclusion, we demonstrate that accurate CV estimates can be inferred in axon populations across a range of configurations, except in some exceptional cases or where axonal density is low. As a proof of concept, for the first time, we generated an *in vivo* map of conduction velocity in the human corpus callosum with estimates consistent with values previously reported from invasive electrophysiology in primates.

## 1 Introduction

The conduction velocity (CV) of action potentials along axons is a key neurophysiological property upon which neural communication depends. While *in vivo* CV measurements in peripheral nerves are comparatively trivial, it is currently not possible to obtain *in vivo* measurements of CV in the central nervous system (CNS). The ability to make such measurements, however, would yield a great deal of insight into how the brain encodes and integrates information and how such mechanisms are optimised in the human brain [1, 2, 3, 4, 5, 6, 7, 8]. Furthermore, being able to image CV in CNS axons *in vivo* would allow us to identify individual differences in CV, and examine how and why CV is altered in healthy development, ageing and disease states.

Previously, simple relationships between axon morphology and CV have been derived from early electrophysiological and theoretical literature [9, 10, 11, 12, 13, 14, 15] (see [16] for a review). In particular, Rushton [12] derived a very simple model to predict conduction velocity from g-ratio and axon diameter. An alternative model derived by Waxman and Bennett [17] models CV as a simple correlation with the outer fibre diameter.

Recent developments in MRI acquisition technology and modelling claim to provide non-invasive estimates of microstructural attributes relevant to CV, including axon diameter (AD) [18, 19], axonal volume fractions (AVF) [20, 21], myelin volume fractions (MVF) [22, 23] and g-ratios [24, 25]. It is tempting, therefore, to speculate that one might use this information to obtain individual specific estimates of CV *in vivo*. To the authors knowledge, no peer-reviewed studies have estimated CV from *in vivo* human MRI, although one study by Horowitz at al [26] has shown correlation between MRI-based estimates of AD and inter-hemispheric transfer delay in electroencephalography, implying MRI-derived measures of AD correlates with CV.

However, beyond the parameters mentioned above, CV depends, to greater or lesser extent, on a number of parameters that are not currently accessible *in vivo*, and yet contribute considerable variability across fibre populations and across individuals. These include the distance between the nodes of Ranvier, inter-nodal spacing, and electrical properties of the axonal and myelin membranes. We address these issues through simulation and then present some CV estimates in human corpus callosum obtained from *in vivo* MRI data

## 2 Sensitivity of CV to axonal parameters

This section addresses the first issue: How sensitive is CV to axonal parameters and are simplified models of CV sufficient to capture variance in CV?

The physiological mechanisms of axon potential propagation have a complex dependency on many parameters that cannot be quantified *in vivo*. In particular, microstructural properties of the nodes of Ranvier, including their length and diameter, contribute to the surface area on which permeable ion channels can reside, impacting on the electrical properties of the axon. Moreover, the inter-nodal distance is important in determining how many instances of depolarisation are required for an action potential to traverse a unit length of axon. Given these various factors, it is important to establish whether it is feasible to obtain accurate estimates of CV from a simplified model using only parameters that have previously been reported as quantifiable using MRI.

A sensitivity analysis on parameters affecting CV has previously been performed [15]. However, this utilised a simple one-at-a-time (OAAT) analysis (where each parameter is varied one at a time) which does not consider *combinations* of parameters, and how interactions between parameter changes affect CV. Moreover, a number of important properties that affect the excitation of the axonal membrane, such as the peri-axonal space, were omitted in that previous analysis. Here, we perform a more comprehensive analysis. We perform extensive simulations of axon physiology using the model of [27] and perform sensitivity analysis to determine the sensitivity of CV across a wide region of the parameter space, and to quantify the variance in CV accounted for by each parameter.

### 2.1 Method

#### 2.1.1 Simulations

The ‘Model C’ axon model of Richardson et al [27], as implemented by [28] (code obtained from https://github.com/AttwellLab/MyelinatedAxonModel) was used to analyse the sensitivity of CV to variance in each of the 14 parameters listed in the upper part of Table 1. Model parameters derived from optic nerve [28, 29]. were used as a proxy for CNS axons. Some parameters were assumed to be well-constrained across individuals and fibre populations and thus not tested (fixed parameters listed in Table 1). Others, such as the number of myelin wraps and myelin thickness, are dependent on g-ratio, AD and myelin periodicity, and so were not directly manipulated. The simulated axon was comprised of 50 laminated inter-nodal regions. All parameters were kept constant across all nodes and inter nodes along the length of the axon. Some internode parameters, such as periaxonal width, were varied at the paranode to accommodate unique morphological characteristics in these regions [30].

**Table 1:** Baseline and range values for each parameter tested for parameters of interest and values for fixed parameters of the Richardson model. All baseline values were those used in [28]. References indicate where values for range were obtained from, otherwise 20% of the baseline value was used

| Varied parameters |  |  |  |
| --- | --- | --- | --- |
| Parameter | Units | Baseline value ($\Phi_i$) [28] | ± Limits ($\vert \Delta\Phi_i\vert $) |
| Axon diameter | $\mu\text{m}$ | 0.82 | 0.31[28] |
| d-ratio (node diameter / axon diameter) | - | 0.9 | 0.18[29] |
| Node length | $\mu\text{m}$ | 1.02 | 0.15[28] |
| Internode length | $\mu\text{m}$ | 139.26 | 31.53[28] |
| Peri-axonal width | nm | 15 | 3 |
| Myelin periodicity | nm | 15.6 | 3.12 |
| g-ratio (inner diameter / outer diameter) | - | 0.78 | 0.057[29] |
| Internode leakage conductance | $\text{mS cm}^{-2}$ | 0.1 | 0.02 |
| Intra-axonal resistivity | $\Omega \text{ m}$ | 0.7 | 0.14 |
| Peri-axonal resistivity | $\Omega \text{ m}$ | 0.7 | 0.14 |
| Myelin conductance | $\text{mS cm}^{-2}$ | 1 | 0.2 |
| Fast Na+ conductance | $\text{mS cm}^{-2}$ | 30 | 6 |
| Persistent Na+ conductance | $\text{mS cm}^{-2}$ | 0.05 | 0.01 |
| Slow K+ conductance | $\text{mS cm}^{-2}$ | 0.8 | 0.16 |
| Parameters dependent on varied parameters |  |  |  |
| Parameter | Units | Baseline value |  |
| Myelin width | $\mu\text{m}$ | 0.101 | |
| Myelin periodicity | nm | 15.6 |  |
| # of myelin wraps | - | 7 |  |
| Node diameter | $\mu\text{m}$ | 0.73 | |
| Internode diameter | $\mu\text{m}$ | 1.05 | |
| Paranode diameter | $\mu\text{m}$ | 1.02 | |
| Paranode length | $\mu\text{m}$ | 2.11 | |
| Periaxonal width at internode | nm | 15 |  |
| Effective periaxonal width at paranode | nm | 0.0077 |  |
| Fixed parameters |  |  |  |
| Temperature | $^{\circ}\text{C}$ | 37 | |
| Number of nodes | - | 50 |  |
| Stimulus amplitude (baseline condition) | nA | 2.73 |  |
| Stimulus duration | ms | 10 |  |
| Axon capacitance | $\mu\text{F cm}^{-2}$ | 0.9 | |
| Node capacitance | $\mu\text{F cm}^{-2}$ | 0.9 | |
| Myelin membrane capacitance | $\mu\text{F cm}^{-2}$ | 0.9 | |
| Node Resting potential | mV | -82 |  |
| Node Reversal potential | mV | -83.4 |  |
| Na+ reversal potential | mV | 50 |  |
| K+ reversal potential | mV | -84 |  |

In the simulations, each model axon was subjected to a current stimulation applied for 10 s to the first node. The amount of current was calibrated such that it produced a peak membrane depolarisation of +50mV in the first node (in the baseline condition, this results in a stimulus amplitude of 2.73 nA). The resultant CV was then measured over a 10-node interval between the 30th and 40th node, except in cases where the CV was too slow for action potentials to reach the 40th node in the simulation duration, in which case the recording interval was moved to earlier segments so that CVs could be measured. To establish that action potentials propagated consistently along the length of the axon, simulations were checked to ensure membrane potential peaks of at least −40 mV were achieved on a minimum of 10 consecutive nodes.

#### 2.1.2 Sensitivity Analysis

Sensitivity was assessed by sampling the corners of a 14-dimensional hypercube in the parameter space, i.e., for every possible combination of positive and negatives changes in each parameter. The dimensions of the hypercube were set to 1 s.d. around the baseline condition (with baseline being the same conditions used for the simulations in [28], given in Table 1), where s.d. was determined from experimental observations in optic nerve [28, 29], or 20% where no such data were available. An exhaustive analysis of 2^14^ = 16, 384 comparisons were made. All simulations ran produced generated action potentials that propagated along the length of the axon.

A one-at-a-time (OAAT) sensitivity analysis was performed for each parameter at 10 equally-spaced intervals within a 20% range around the baseline condition (see B). This shows that relative changes in CV are approximately linear with change in parameter so we can assume sampling only the corners of the hypercube is sufficient to capture the variability in CV within this region of the parameter space. The change in CV, 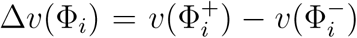, due to a change in each parameter 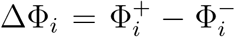, relative to the CV of the baseline condition 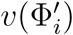 was calculated. The corresponding sensitivity was calculated by normalising the relative change in CV to the relative change in the parameter.

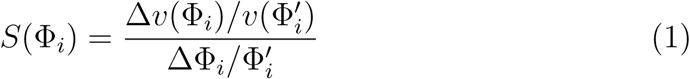

#### 2.1.3 Testing Simplified Models of CV

We aimed to derive a simple model to predict CV (and associated variance) from the two parameters that have previously been reported as accessible from in vivo MRI: g-ratio, *g*, and internal AD, *d*. We tested the model across a grid comprising 10 equally-spaced values of *d* (0.25 to 8 µm) and 12 equally-spaced values of *g* (0.4 to 0.95). For each grid-point, we repeated the sensitivity hypercube analysis by running the Model C [27] simulation across all possible combinations of the remaining non-MRI accessible parameters, to generate a distribution of CVs for each point on the grid. This resulted in 10 × 12 × 2^12^ = 491, 520 model runs. The mean and standard deviation of CV at each point was calculated We then fitted simplified models based on the Rushton formula [12] and the linear relationship with outer diameter [17]. We also explored some more complex polynomial models that could potentially provide better fits to the data. In all cases, metrics of the model fit performance and parsimony, including Akaike and Bayesian information criterion (AIC and BIC) were computed. As with the sensitivity analysis, simulations were checked to ensure that action potentials were successfully generated and propagated along the axon.

### 2.2 Results

The conduction velocity obtained in the baseline condition was 2.95 ms^−1^, in agreement with the original simulations in [28] (see also A for further validation). Action potential propagation was successful in all simulations. The distribution of relative changes in CV, due to change in each parameter, is shown in Figure 1(a), while Figure 1(b) shows the total variances in CV due to change in each parameter relative to the total variance. The majority of the variance is explained by AD, followed by internode length and then g-ratio. A key finding of this analysis is that combined together, AD and g-ratio explain 89.2% of the model variance in CV.

**Figure 1:**
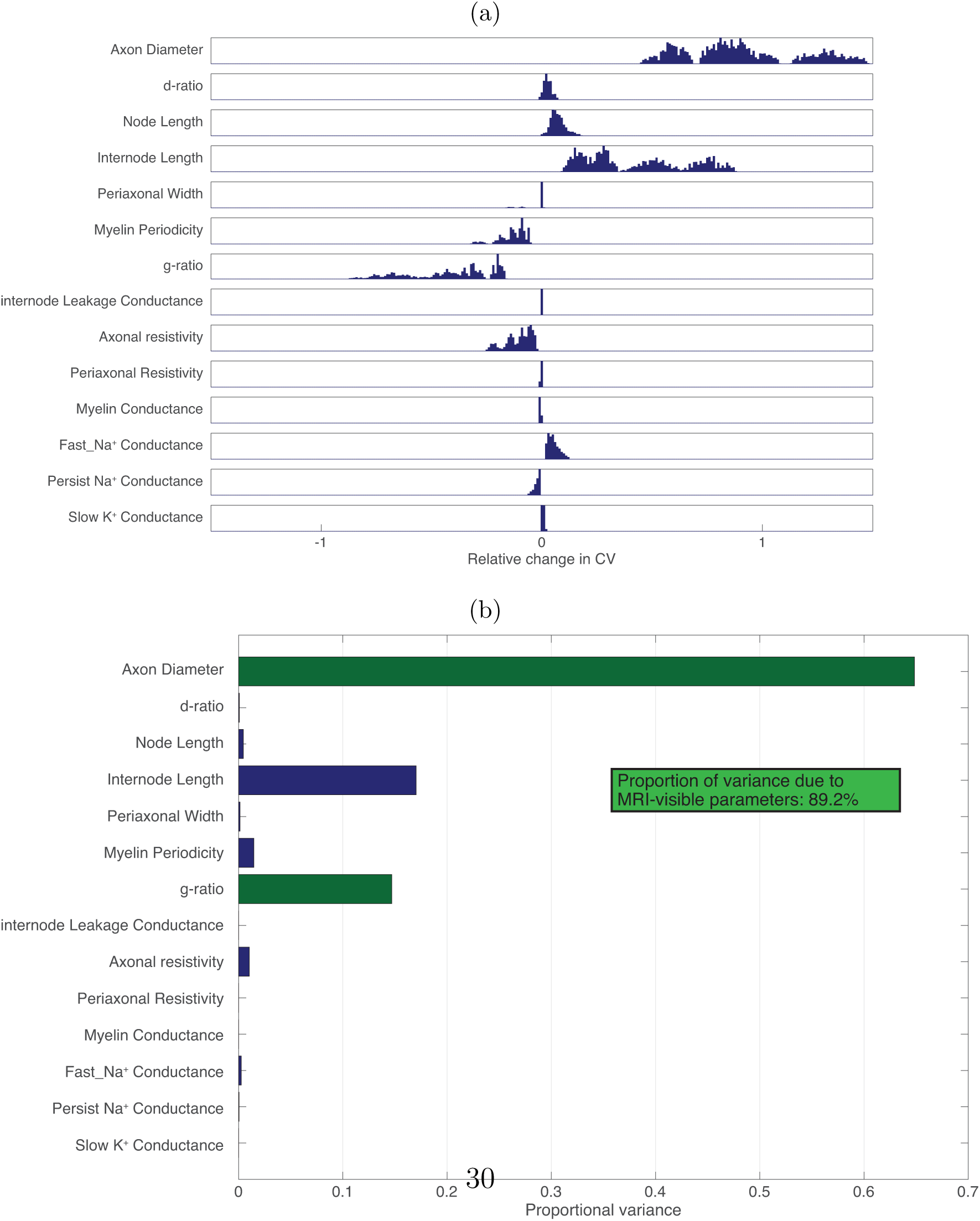
(a) Distributions of proportional change in CV for a stepwise change in each parameter (parameter step size determined by limits indicated in Table 1) across all points in the parameter space. (b) The total variance for each parameter step change as proportion of variance across all simulations. MRI-visible parameters indicated by green bars.

The distribution of relative sensitivities of CV to unit changes in each parameter are shown in Figure 2(a) while Figure 2(b) shows the sum-squared sensitivity for each parameter, proportional to the sum-squared sensitivity across all parameters. CV is most sensitive to unit change in *g* by a considerable margin. *d* has the second highest sensitivity. Combined together, *d* and *g* account for 91.5% of the total sensitivity of CV.

**Figure 2:**
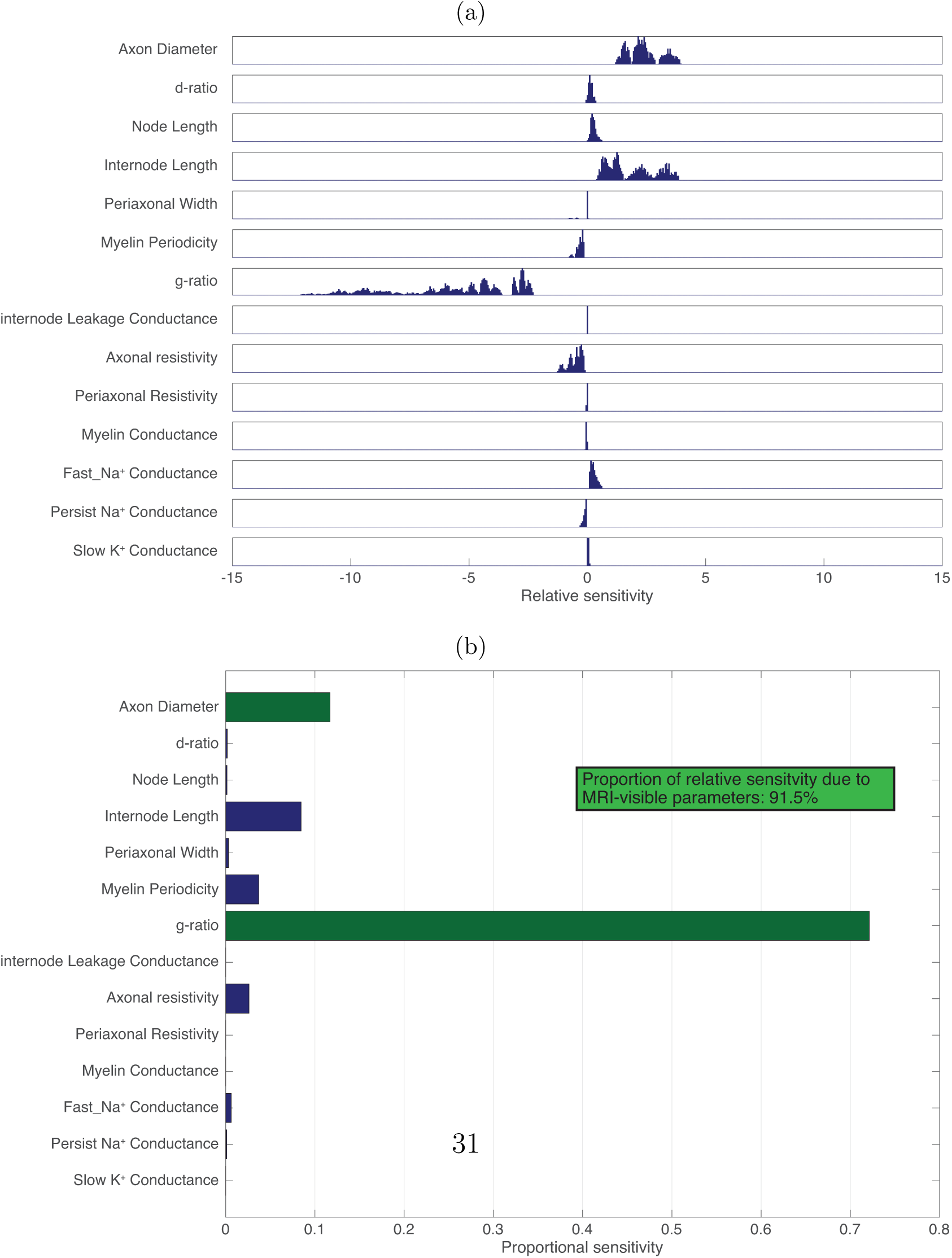
(a) Distributions of relative sensitivities of CV to unit change in each parameter across all points in the parameter space. (b) The sum-squared relative sensitivity for each parameter step change as proportion of the total sum-squared sensitivity across all simulations. MRI-visible parameters indicated by green bars.

The distribution of CVs across *d* and *g* are shown in Figure 3. 7 grid points tested failed to produce action potentials (where AD is 0.25µm and g-ratio is above 0.65) as indicated by missing points in Figure 3. The mapping of CV to *d* appears to approximately linear, while the mapping to *g* follows an inverse log square root function. This is similar to the form given by [12].

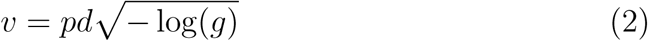

where *p* is some constant of proportionality, fitted to *p* = 6.644 (confidence bounds: [6.638, 6.651]). The 2D fitting to the original Rushton model yielded a good fit (SSE=1.54 × 10^7^, *R*^2^ = 0.766), but the fit was poor where *d* is large and *g* is small (Figure 4).

**Figure 3:**
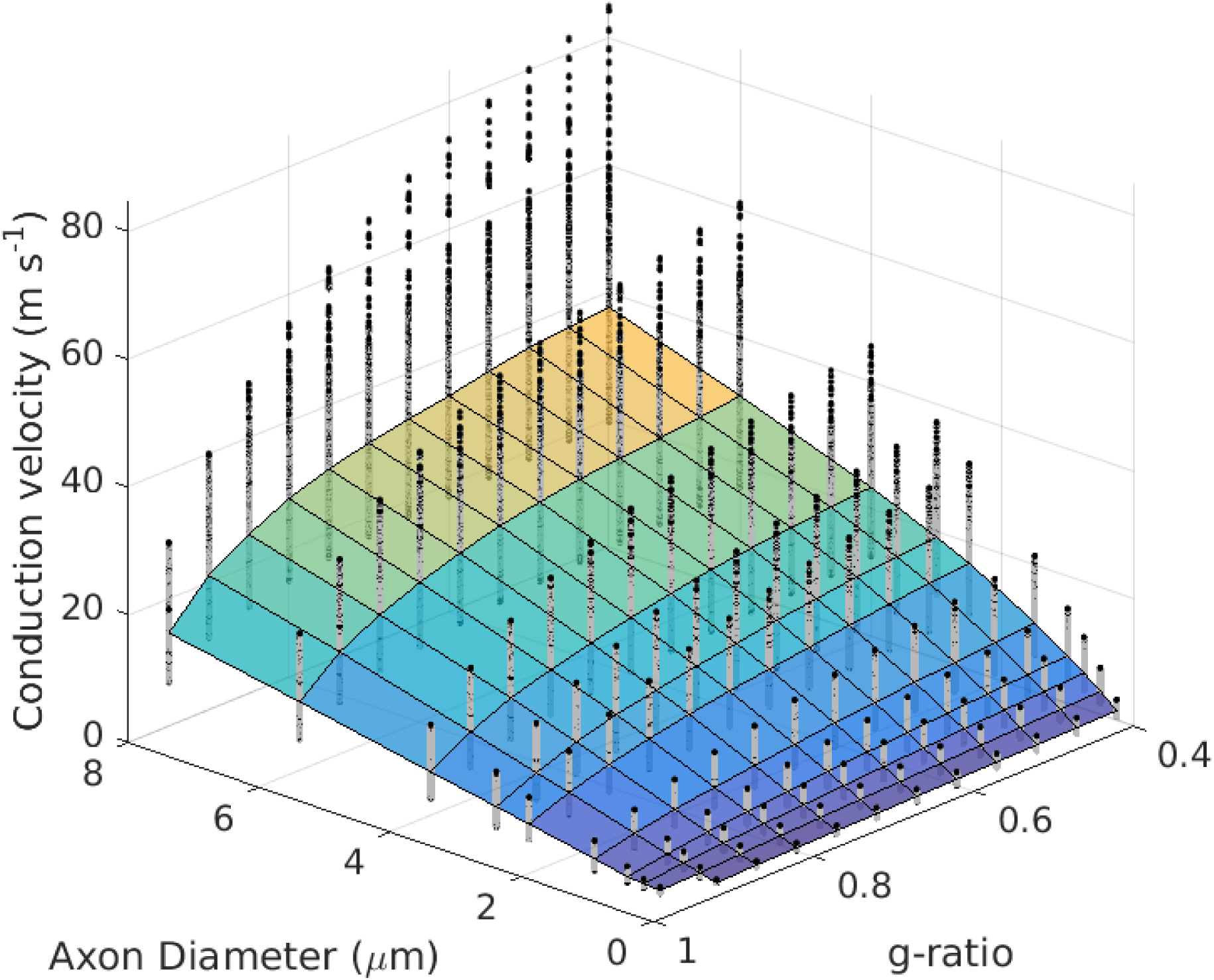
Distribution of CV estimates across fixed values of AD and g-ratio. Surface plot indicates the mean value. Black dots show the distribution of CV estimates at each point.

**Figure 4:**
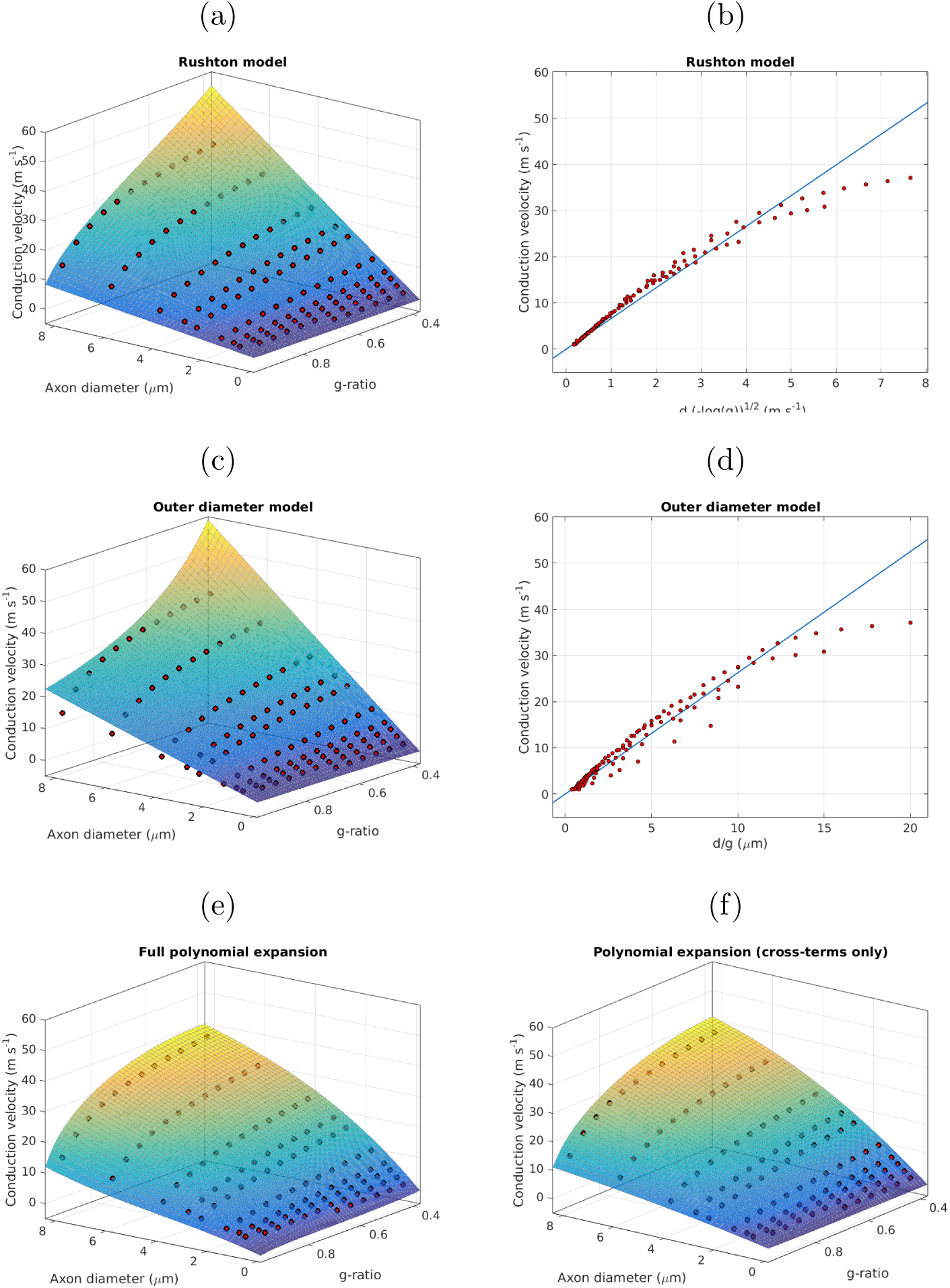
Plots of simplified models fitted to simulated data points across AD, (*d*) and g-ratio (*g*) values.(mean for each AD-g pair indicated by red circles). (a) Rushton model as fitted across values of *d* and *g*; (b) Rushton model as a linear fit to 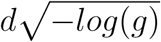 (c) outer diameter model as fitted across values of *d* and *g*; (d) outer diameter model as a linear fit to outer diameter (*d/g*); (e) Full 3rd order polynomial expansion in *d* and 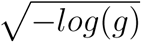 (f) the same polynomial expansion only considering the cross-terms.

We also tested whether CV could be predicted from a linear function of outer diameter [17]. This is simpler to calculate since it uses only one parameter but implicitly assumes a constant *g*-ratio.

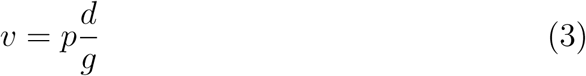

where *p* was fitted to *p* = 2.626 (confidence bounds: [2.624, 2.629]). The goodness of fit was slightly poorer for this model compared to the Rushton model (SSE=1.58 × 10^7^, *R*^2^ = 0.760). The AIC and BIC were comparable to the Rushton model.

Further comparison was made between the two models by computing the SSE for each *d*-*g* pair and plotting the difference in SSE (Figure 5). This shows that where AD is high (above 5 µm), there is a better fit (lower SSE) for the Rushton model where *g* lies between 0.5 and 0.75. The outer diameter model shows better fit where *g* lies between 0.75 and 0.95.

**Figure 5:**
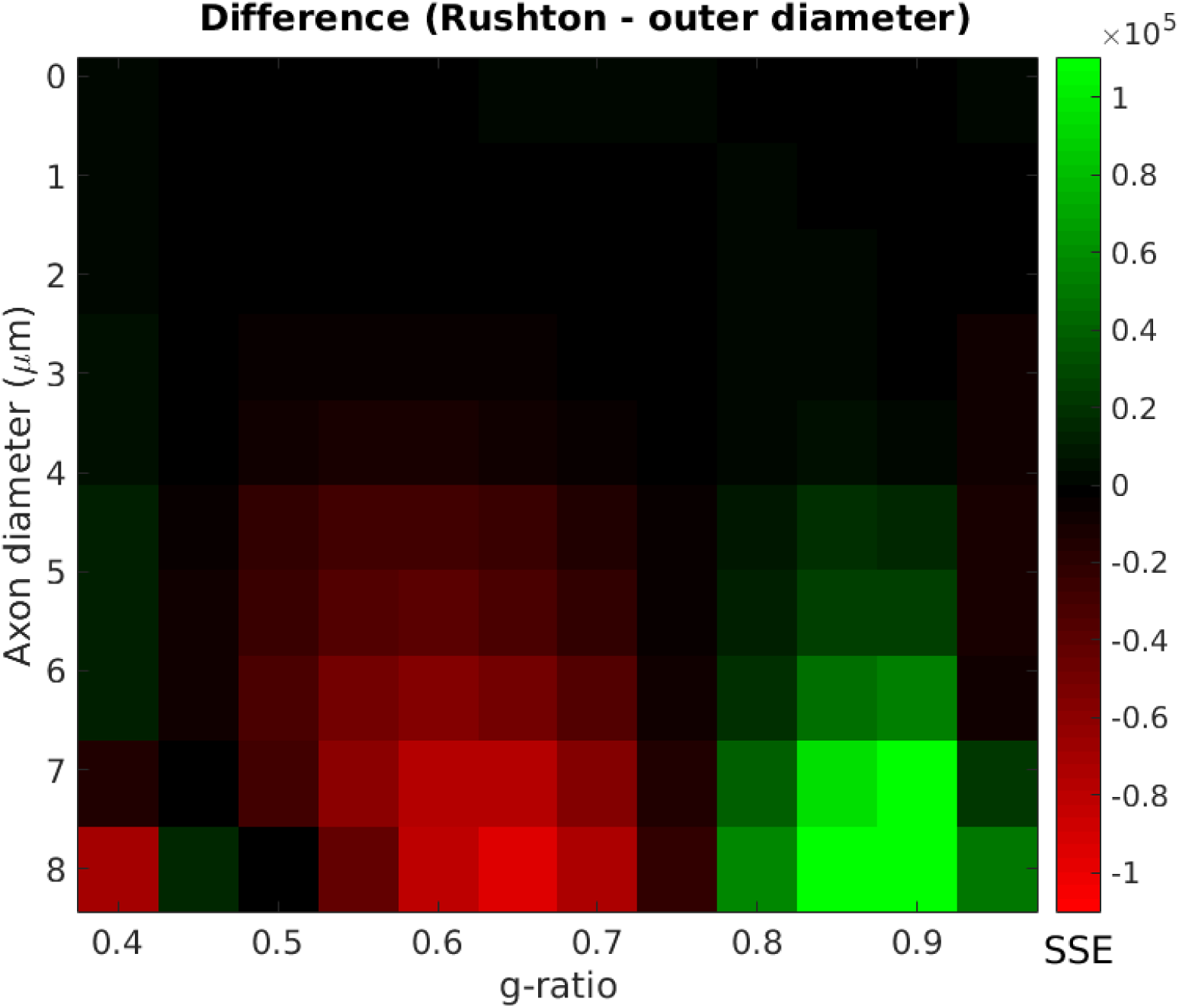
Difference in SSE between the Rushton and linear outer-diameter models. Positive values (green) show higher SSE for The Rushton model, negative values (red) show higher SSE for the outer diameter model.

Two more complex models were tested to compare with the Rushton and linear outer diameter models. A 3rd order 2D polynomial expression in *d* and 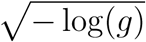 yielded a better fit (SSE=1.23 × 10^7^,*R*^2^ = 0.813) but required fitting of 11 coefficients. A good fit was also achieved when considering only cross-terms in the polynomial, (SSE=1.25 × 10^7^,*R*^2^ = 0.810) which only requires 3 coefficients.

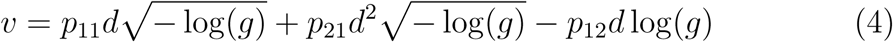

However, the AIC and BIC are lowest for the Rushton model. Therefore, this remains the preferred model for predicting CV (Figure 4). The s.d. of the modelled CVs scaled linearly with the mean CV (SSE=7.22,*R*^2^ = 0.994).

## 3 Estimating CV from MRI-derived parameters

The second part of this study focuses on the second issue highlighted in the introduction: Is it possible to obtain accurate CV estimates from parameters derived from existing microstructural MRI techniques [18, 19, 20, 21, 22, 23, 24, 25]?

All these techniques work by fitting microstructural parameters to biophysical models of the MRI signal using some numerical optimization routine. This approach has some inherent issues. MRI signals are subject to noise from a range of sources. There are problems with fitting model parameters to MRI signals, including degeneracy of solutions in the optimisation process, and the likelihood of fitting the model to noise contributions. As a result, there can be considerable bias in MRI-derived microstructural metrics [31]. We note, in particular, that quantification of inner AD is challenging, if not impossible, at gradient strengths found on typical clinical MRI system (up to 80 mT/m) [32, 33, 34, 35]. This was a criticism levied at the study of Horrowitz et al [36, 33]. However, the advent of ultra strong gradient systems (300 mT/m) provides sensitivity to axon diameter, at least over a limited but relevant range (i.e. above 3 µm) [34]. In this work we therefore focus on simulation (and real data) on an ultra strong gradient system. Although this is a special case, it does allow us to evaluate the feasibility of estimating CV *in vivo*.

This issue of model bias can become an even more pernicious if some models take as input the output of other models, leading to propagation of noise and bias through different models. It is imperative, therefore, that MRI-derived estimates of CV are robust to such errors, which is the subject of investigation in the present study.

### 3.1 Method

To model the effects of MRI noise, MRI data were simulated using analytical expressions for three biophysical models, the Composite Hindered and Restricted Model of Diffusion (CHARMED) [20], the AxCaliber model [18] and multicomponent driven equilibrium single pulse observation of T1/T2 (mcDESPOT) relaxometry [22].

### 3.2 Core biophysical simulations

A single population of axons with a Poisson distribution of diameters (mean AD is parameterised by λ) was simulated with dispersion or crossing-fibre configurations were simulated. The biophysical parameters of the system are listed in Table 4. Systems with this configuration were simulated for a range of AVFs, axon diameters and g-ratios. The g-ratio value is treated as an aggregate measure of g-ratio across the volume. The value AVF ranged from 0.05 to 0.4; mean ADs from 0.25 to 12 µm and g-ratios of 0.4 to 0.9.

**Table 4:** Fixed biophysical parameters used for the MRI simulations

| **Fixed biophysical parameters** |  |  |
| --- | --- | --- |
| Parameter | Units | Value |
| Intracellular axial diffusivity | $\text{cm}^2 \text{ s}^{-1}$ | 2.9 |
| Extracellular axial diffusivity | $\text{cm}^2 \text{ s}^{-1}$ | 1.3 |
| Extracellular radial diffusivity | $\text{cm}^2 \text{ s}^{-1}$ | 0.3 |
| Orientation ($\theta$) | rad | $\pi / 2$ |
| Orientation ($\phi$) | rad | 0 |
| Myelin $T_1$ | ms | 465 |
| Myelin $T_2$ | ms | 26 |
| Myelin residence time | ms | 180 |
| Extracellular $T_1$ | ms | 1070 |
| Extracellular $T_2$ | ms | 50 |
| CSF $T_1$ | ms | 4000 |
| CSF $T_2$ | ms | 2500 |
| CSF volume fraction |  | 0.05 |

#### 3.2.1 Diffusion MRI simulation

CHARMED and AxCaliber MRI data were simulated in MATLAB using parameters that matched a standard protocol used on a Siemens 300 mT/s Connectom system (Table 3). The CHARMED model was then fitted to the simulated data using particle swarm global optimization to handle multiple local minima in the optimisation landscape [37].

**Table 3:** Acquisition parameters used for simulations of diffusion and relaxometry MRI data and for *in vivo* data acquisition

| Parameter | Value |
| --- | --- |
| **Diffusion acquisitions** |  |
| Flip angle | 90° |
| Slice thickness | 2 mm |
| Field of View | 220 × 220 mm |
| Matrix size | 110 × 110 |
| Voxel size | 2 × 2 × 2 mm |
| CHARMED |  |
| $b$ | [500,1200,2400,4000,6000] s mm<sup>-2</sup> |
| # directions | [30,60,60,60,60] |
| $\delta$ | 7 ms |
| $\Delta$ | 23.3 ms |
| Echo time | 48 ms |
| Repetition time | 2600 ms |
| AxCaliber |  |
| $b$ | [2000,4000] S mm<sup>-2</sup> |
| # directions | [30,60] |
| $\delta$ | 7 ms |
| $\Delta$ | [17.3,30,42,55] ms |
| Echo time | 80 ms |
| Repetition time | 3900 ms |
| **Relaxometry acquisitions** |  |
| Slice thickness | 1.72 mm |
| Field of View | 220 × 220 mm |
| Matrix size | 128 × 128 |
| Voxel size | 1.72 × 1.72 × 1.72 |
| SPGR |  |
| Flip angles | [3,4,5,6,7.5,9,12,15,18] ° |
| Echo time | 1.9 ms |
| Repetition time | 4.2 ms |
| IR-SPGR |  |
| Flip angle | 5 ° |
| Echo time | 1.9 ms |
| Repetition time | 4.2 ms |
| Inversion Time | 450 ms |
| SSFP |  |
| Flip angles | [10,10,15,15,20,20,30,30,40,40,50,50,60,60] ° |
| Phase cycle angles | [0,180,90,270,0,180,90,270,0,180,90,270,0,180,90,270] ° |
| Echo time | 2.27 ms |
| Repetition time | 4.54 ms |
| Slice thickness | 1.72 mm |

#### 3.2.2 relaxometry MRI simulation

mcDESPOT MRI data were simulated using the ‘qisignal’ function in the Quantitative Imaging Toolbox (QUIT) [38]. The protocol comprised 8 spoiled gradient recalled (SPGR) images with varying flip angles and 16 steady-state free precession (SSFP) (Table 3) images distributed across 8 flip angles and 4 phase cycle angles. To account for the influence of radio frequency field strength (*B*_1_) and off-resonance frequency (*F*_0_) in the fitting, a range of *B*_1_ and *F*_0_ values were simulated for each noise measurement. To replicate the noise profile obtained from SNR measurements across flip angles, noiseless data were simulated and Rician noise with flip-angle-specific s.d. was added to the simulated data (see C). A 3-pool model (modelling contributions from myelin, extra-cellular and CSF water) was then fitted to the simulated data using the ‘qimcdespot’ function in the QUIT toolbox.

Since mcDESPOT gives a myelin water fraction (MWF) map, as opposed to a true MVF, we estimated the true MVF from the formula:

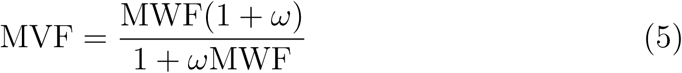

where *ω* = 1.44 is the ratio of lipid to water in the myelin [39] (see D for derivation).

#### 3.2.3 g-ratio and CV estimation

g-ratios were computed using the approach of Stikov et al [24]:

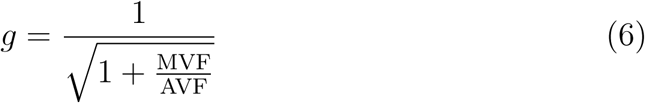

This approach has been shown to give a valid aggregate measure of g-ratio for a distribution of ADs. Using the Rushton model (Eq. 2), with *p* = 6.65 as fitted previously using simulations, CV was estimated for each combination of mean AD and g-ratio.

#### 3.2.4 Noise

Noise was simulated by adding Rician noise to each simulated MRI acquisition. The s.d. for each acquisition was modified to replicate the SNR profiles observed in real data (see C). Additionally, to test sensitivity to noise, data simulations were repeated with noise s.d. at 50% and 200% of the original noise s.d. This was done for all permutations across the 3 MRI parameters. For all simulated acquisition and permutations of noise levels, 100 iterations were performed. This resulted in a total of 100 × (1 + 2^3^) × 11 × 8 = 79, 200 diffusion MRI simulations and 100 × (1 + 2^3^) × 8 × 12 = 86, 400 relaxometry MRI simulations.

#### 3.2.5 Error measurement

Errors in CV estimates were quantified in 4 ways: Bias, variance, relative sensitivity to modelling errors and relative sensitivity to noise. Bias was quantified by the mean relative error in CV, variance was quantified by the variance of the CV estimates normalised to the original CV estimate. Relative sensitivity to modelling errors was quantified by taking the ratio of the relative error in CV to the relative error of the relevant imaging parameter (AVF, AD and MVF). Sensitivity to noise was estimate by taking the difference in CV estimates between the 50% and 200% noise condition and normalising to the difference in noise s.d.

## 3.3 Results

### 3.3.1 Errors in modelled parameters

Errors in relevant fitted parameters are shown in Figure 6 and distributions of errors across parameters are shown in Figure 7. Overall, the lowest errors are in AVF (mean ± s.e.: 0.029 ± 0.0002) with higher errors for MVF (0.48 ± 0.0071) and AD (0.65 ± 0.0072). AVF estimates show the highest error for the smallest values of AVF, and this error decreases with increasing AVF up to value of AVF = 0.5. Across all other parameters (AD, MVF and g-ratio), errors were also largest for the lowest AVF and decreased with increasing AVF. The optimal value for MVF and g-ratio (where the relative error is lowest) scales approximately linearly with the AVF. Errors in AD are largest when AD and AVF are low.

**Figure 6:**
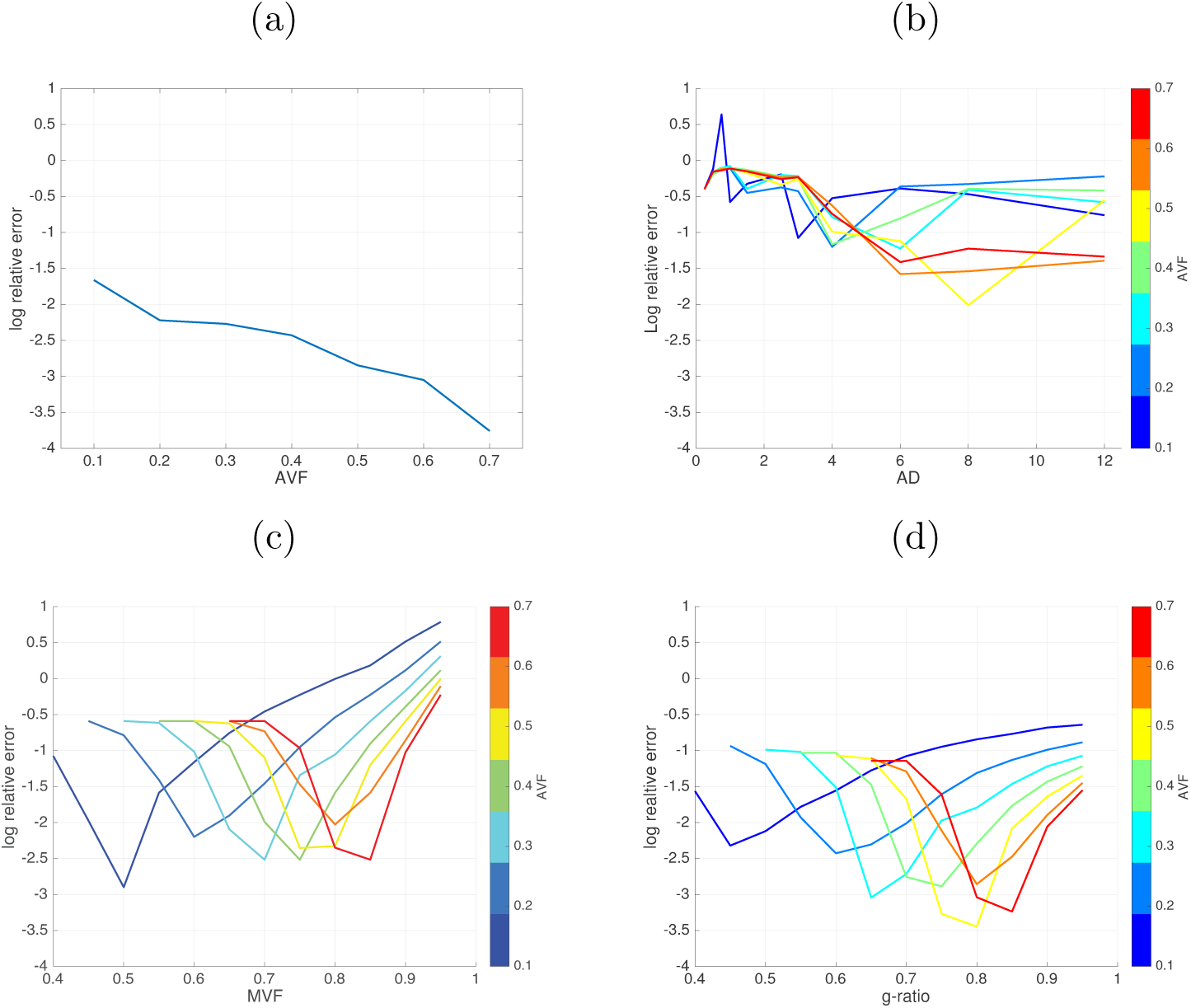
Log relative errors in (a) AVF (b) AD (c) MVF and (d) g-ratio.

**Figure 7:**
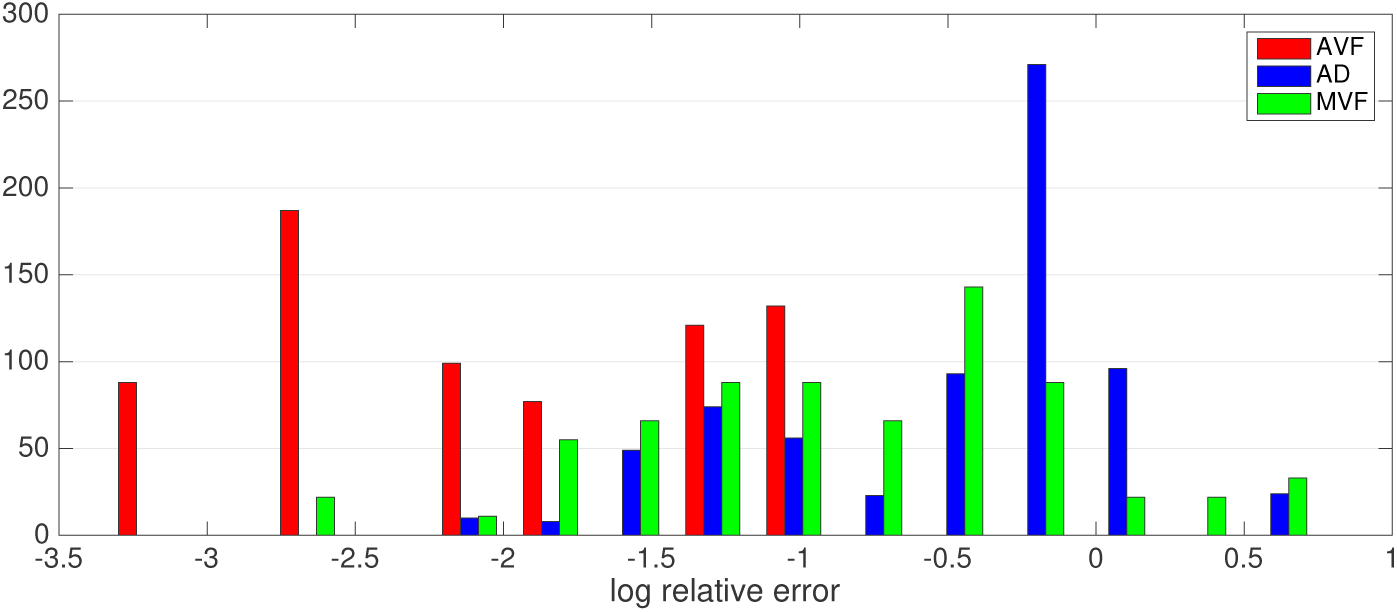
Distributions of log relative errors in AVF (red), AD (blue) and MVF (green) across all iterations and parameters

### 3.3.2 Bias in CV estimates

Relative errors in CV across the parameter space tested are shown in Figure 8. The CV estimates show a less than 5% bias across a region of parameter space where AVF is 0.25 or above, AD is below 10µm. There is little dependency on g-ratio. Bias is greatest (over 50%) in regions where AVF is low (below 0.25) and AD is between 2-4 µm or greater than 8 µm.

**Figure 8:**
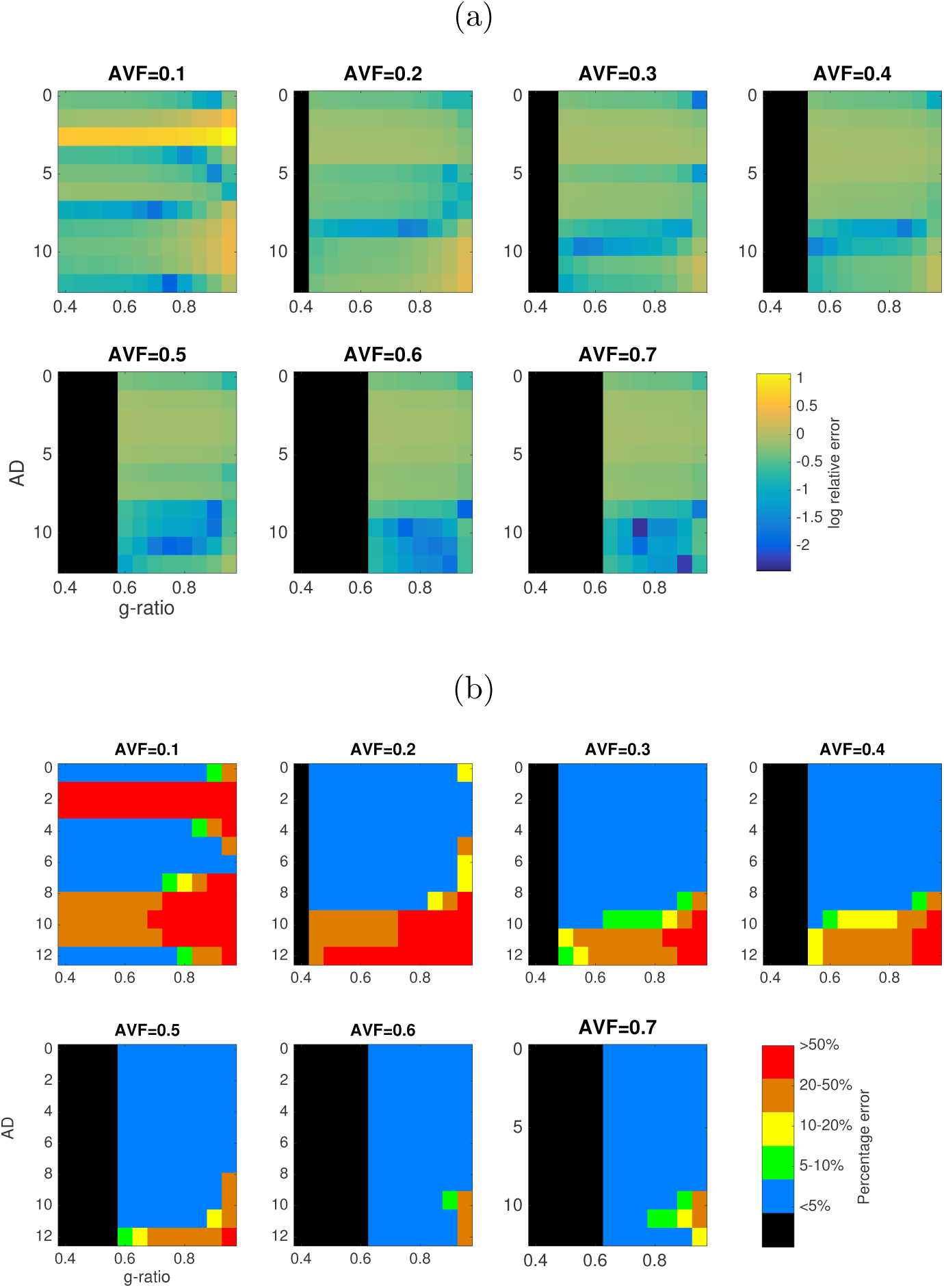
(a) Log relative error in CV estimates across values of AVF, AD and g-ratio. (b) Regions of parameter space where relative variance is less than 5% (blue), 5-10% (green), 10-20% (yellow), 20-50% (orange) and greater than 50% (red) error in CV estimates. Black regions are where axon AVF/g-ratio combination gives an infeasible MVF.

### 3.3.3 Variance in CV estimates

Variance in CV estimates is shown in Figure 9. The normalised variance is below 0.5 where AVF is high (above 0.25) and AD is between 1 and 10 µm.

**Figure 9:**
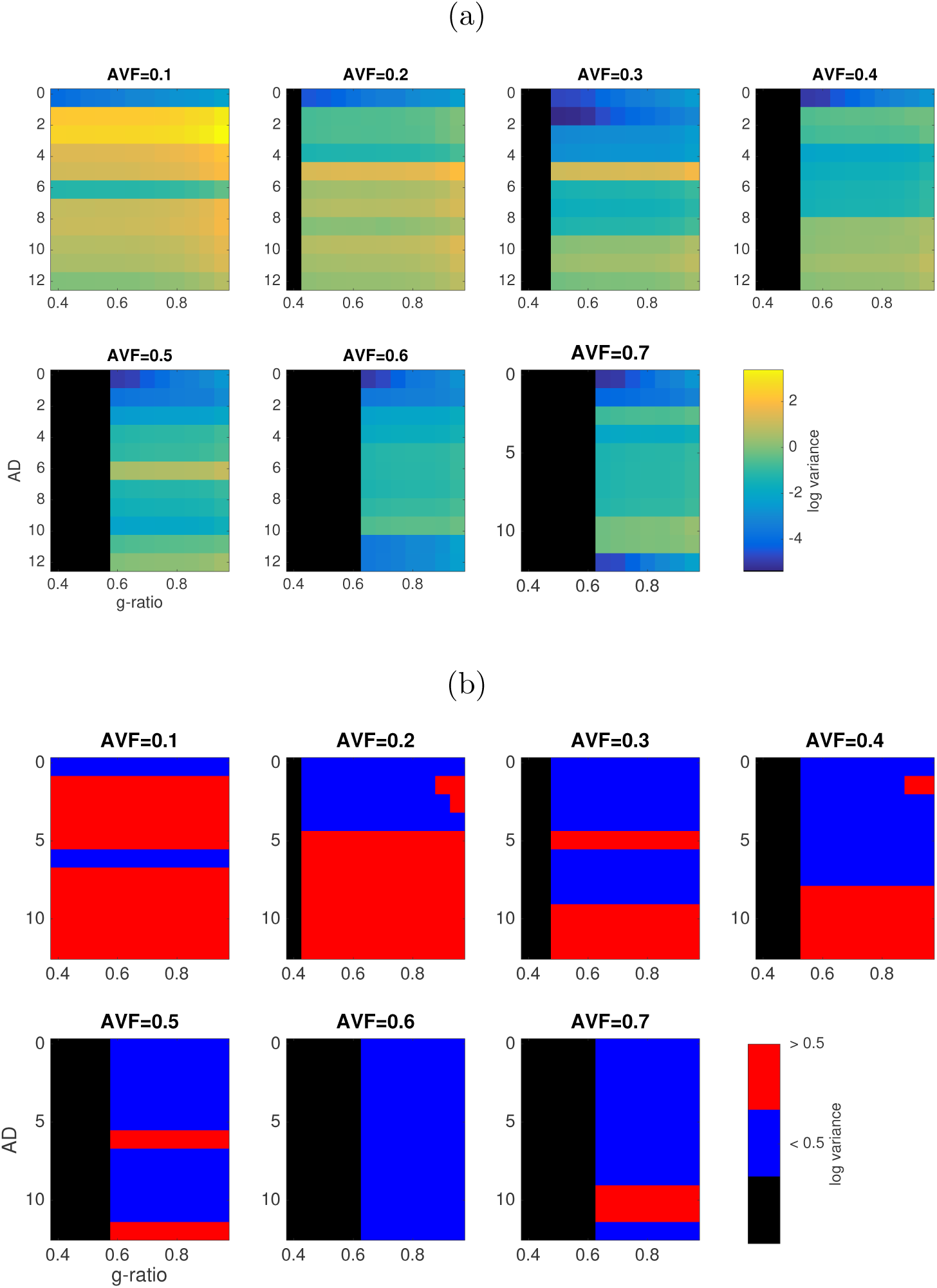
(a) Log normalised variance in CV estimates across values of AVF, AD and g-ratio. (b) Regions of parameter space where normalised variance is less than 0.5 (coloured in blue) or greater than 0.5 (coloured in red).

### 3.3.4 Sensitivity to parameter errors

Distributions of relative sensitivities to modelled parameter errors across the tested parameter space are shown in Figure 10. The sensitivity of CV to errors in AVF (mean ± s.e.: 5.02 ± 0.006) is much higher than to errors in AD (0.39 ± 0.007 µm) and in MVF (2.38 ± 0.014). The proportional variances in CV estimates explained by errors in the three MRI parameters are shown in Figure 11. Errors in AVF derived from CHARMED have the highest effect overall. Errors in MVF derived from mcDESPOT contribute more to variance in regions of low g-ratio (below 0.7) and low AVF (below 0.2). Errors in AD derived from AxCaliber contribute very little to variance in CV across all parameters.

**Figure 10:**
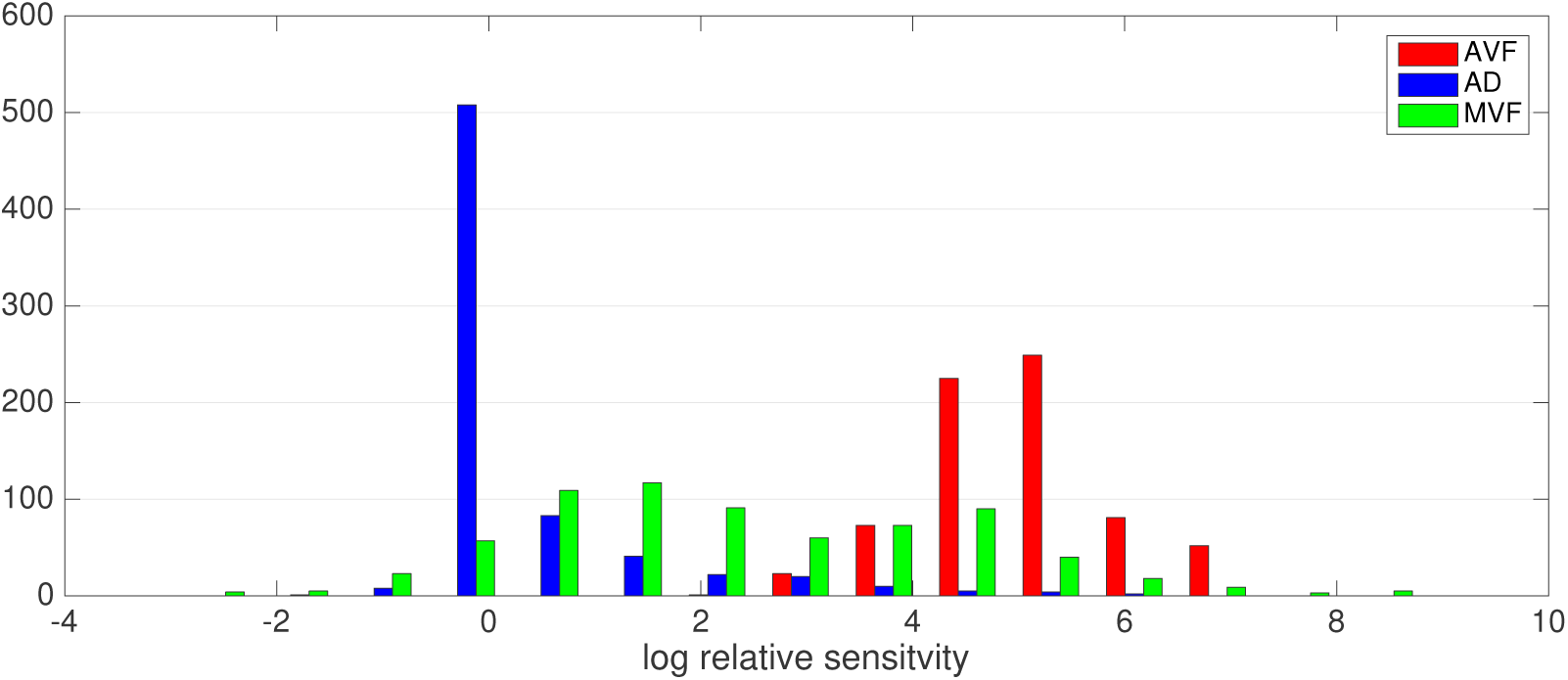
Proportional variance explained by errors in each MRI parameter, across the parameter space tested.

**Figure 11:**
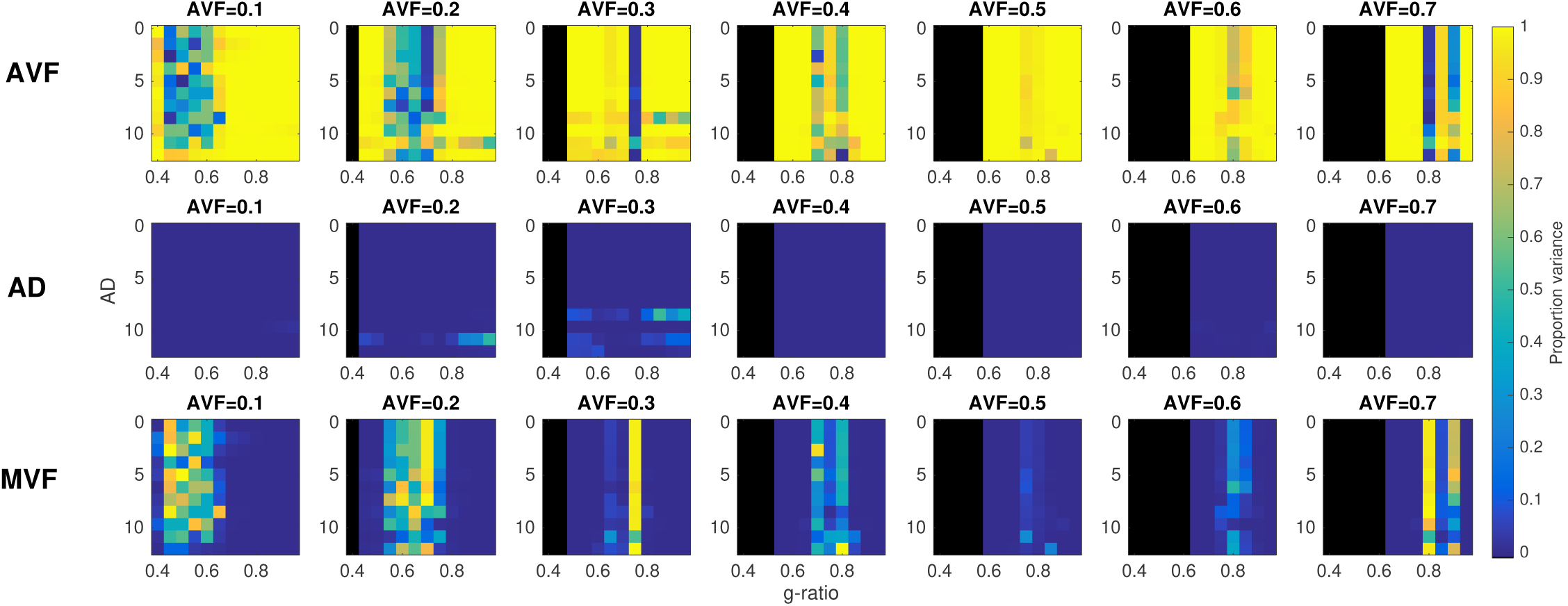
Distributions of log relative sensitivity of CV to errors in AVF (red), AD (blue) and MVF (green) across all iterations and parameters.

### 3.3.5 Sensitivity to MRI noise

Distributions of relative sensitivities to noise across the tested parameter space are shown in Figure 12. The proportional variance in CV estimates explained by noise across three MRI parameters are shown in Figure 13. Proportional variances are quite uniform across the parameter space, with noise in relaxometry acquisitions (MVF) contributing almost nothing to the variance in CV. Noise in diffusion acquisitions (AVF and AD) each contribute equally about 50% of variance in CV.

**Figure 12:**
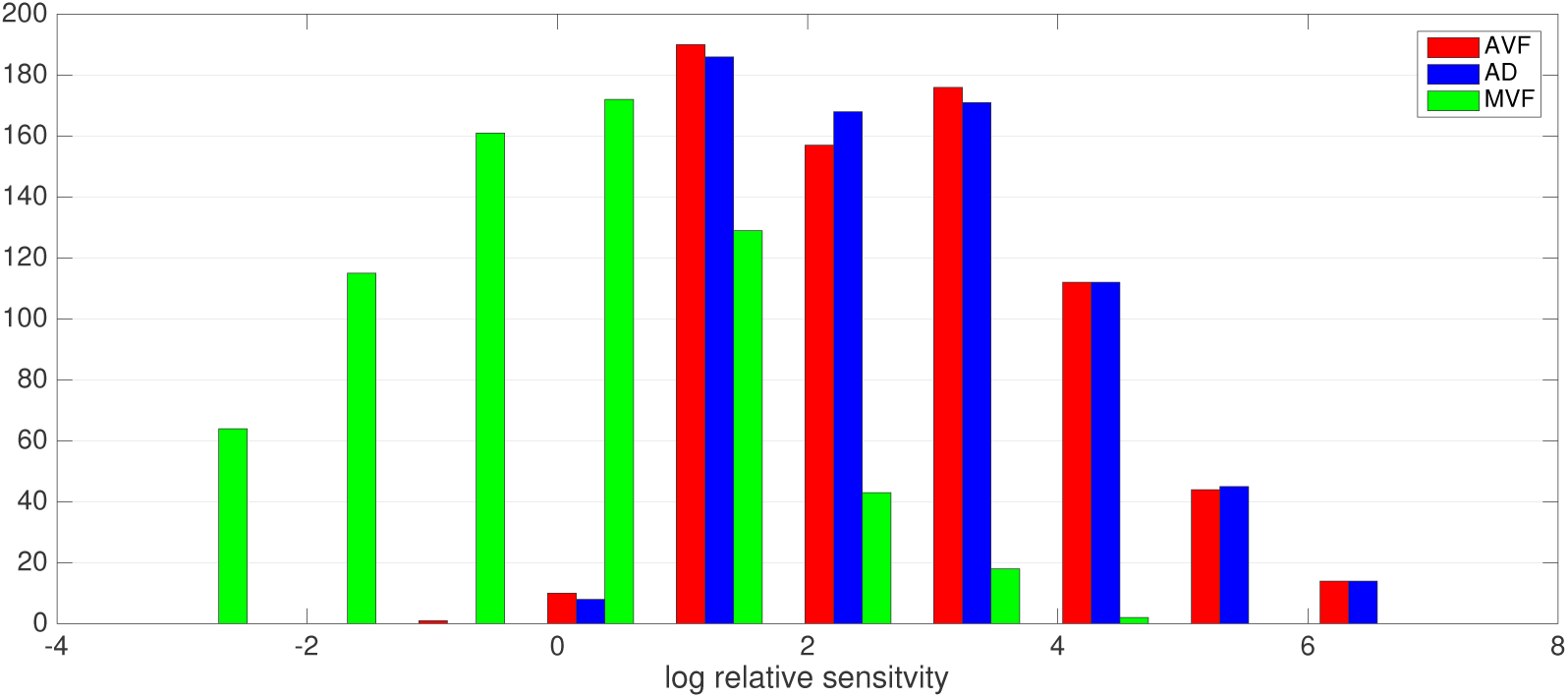
Distributions of log relative sensitivity of CV to noise in AVF (red), AD (blue) and MVF (green) acquisitions across all iterations and parameters.

**Figure 13:**
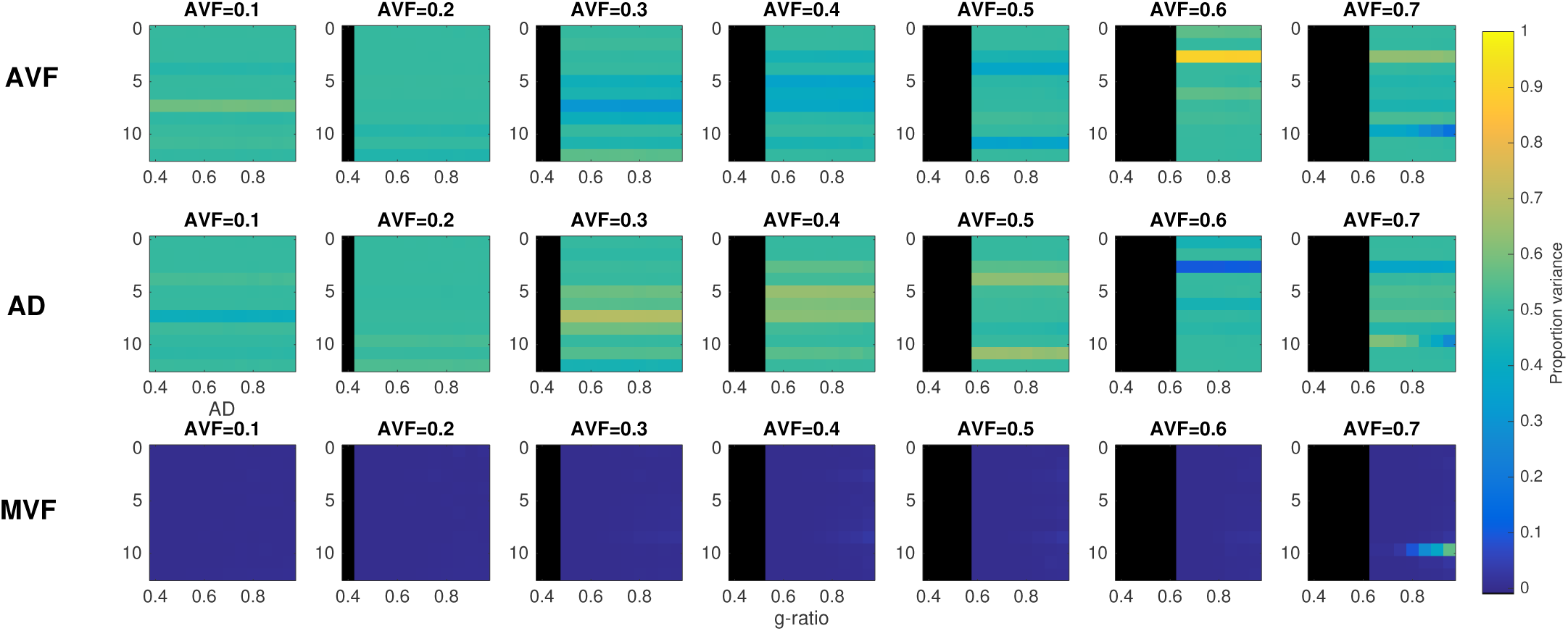
Proportional variance explained by noise in MRI acquisition, across the parameter space tested.

## 4 *in vivo* CV estimates from human MRI data

As a proof of principle we apply the proposed approach to *in vivo* human data obtained from. Subject to the caveats regarding the sensitivity to AD. Data were acquired on a high gradient MRI system. The analysis focuses on the corpus callosum as the axons here have a relative uniform orientation, minimal dispersion and have an AD that is in the range that is detectable on a high-gradient MRI system.

### 4.1 Method

#### 4.1.1 MRI acquisition

CHARMED, AxCaliber and mcDESPOT data were all acquired from a single healthy human participant (F,28y) on a Siemens 3T 300mT/m Connectom system. The acquisition parameters used were identical to those used in the simulations (see Table 3).

#### 4.1.2 Diffusion MRI processing

Motion, eddy current and EPI distortions were corrected using FSL TOPUP and EDDY tools [40]. Gradient non-linearities were corrected. All diffusion data were then registered to a skull-stripped [41] structural T1-weighted image using EPIREG [40]. AVF and AD parameters were fitted to the CHARMED and AxCaliber models using the same approach described for the MRI simulations.

#### 4.1.3 Relaxometry MRI processing

Motion correction was applied to the SPGR and SSFP data using FSL mcFLIRT and then the brain was skull-stripped [41]. All subsequent fitting steps were performed using the QUIT toolbox [38]. A *B*_1_ map was estimated by fitting the data to the DESPOT1-HIFI model [42] and then fitting to a 8th order 3D polynomial. An *F*_0_ map was estimated by fitting to the DESPOT2-FM model [43]. These were then used for the final fitting to the mcDESPOT model, as described for the MRI simulations. The final MVF maps were registered to the T1-weighted image using FLIRT [40] so that all parameter maps were in the same space.

#### 4.1.4 CV mapping

AVF, AD and MVF parameters were fitted using the same methods described for the MRI simulations and CV maps were generated using the same approach. In addition to generating a CV map, maps of estimated bias and variance were obtained by interpolating the estimates obtained from the MRI simulations using the MRI-estimated MVF, AD and g-ratio values. The bias was then used to obtain a bias-corrected CV map.

### 4.2 Results

*In vivo* MRI data in the corpus callosum are shown in Figure 14(a). CV estimates in the corpus callosum are between 8.1 and 41.6 ms^−1^ (median: 14.2 ms^−1^). Smaller CV estimates are seen in the genu and splenium of the corpus callosum, consistent with the fact that these regions have smaller ADs. The bias in these regions is higher than in the body of the corpus callosum. The bias-corrected CV values showed a range of 3.7 - 20.4 ms^−1^ (median: 12.2 ms^−1^). This is very similar to the range observed in Macaque corpus callosum [44] (2.8 - 22.5 ms^−1^, median = 7.4 ms^−1^, see Figure 14(b)).

**Figure 14:**
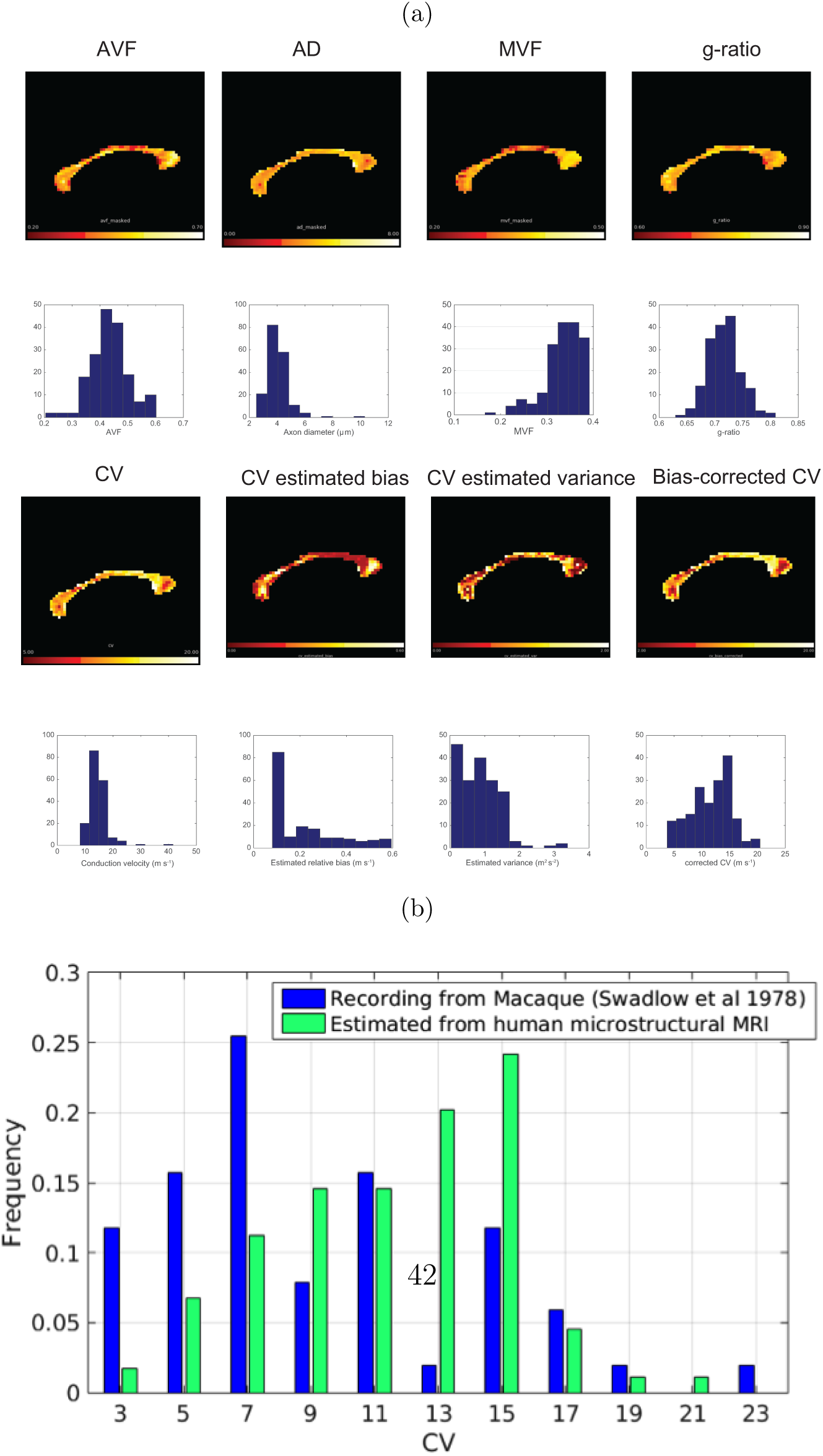
(a) Fitted in vivo human MRI data to microstructural parameters in corpus callosum, and corresponding histograms showing the distribution of values across the corpus callosum (b) comparison of distribution of CV estimates from MRI and recordings from Macaque electrophysiology [44].

## 5 Discussion

This paper has explored the feasibility of obtaining conduction velocity (CV) maps from *in vivo* human MRI, using a simplified model of axonal CV. Results from the axon simulations demonstrate that 89.2% of the variance in CV, and 91.5% of the sum-squared sensitivity of CV, can be attributed to variance in AD and g-ratio. Looking at variance (using ecologically valid variances in parameters where possible), implicate AD as the most important parameter, while looking at sensitivity to a unit change in parameter, g-ratio is implicated as the most important parameter. Therefore, considering the fact that AD varies much more in axon populations than g-ratio, capturing accurate estimates of AD is still more important than g-ratio for CV estimation.

The Rushton and outer diameter models for CV provide a reliable estimate of CV from MRI-derived estimates of g-ratio and AD. In addition, we show that it is possible to account for uncertainty in CV estimates due to parameters not accessible *in vivo*. Thus, when the reliable estimates of AD and g-ratio can be made, it is feasible to obtain estimates of axonal CVs *in vivo*. The match in the parsimony measures (AIC/BIC) for the Rushton model and outer-diameter model (Table 2) were comparable, with a minor improvement in SSE for the Rushton model. Indeed, [45] used the outer-diameter model to good effect (see also [33] for a discussion fo merits of the outer diameter model). However, examining the regional difference in the SSE (Figure 5) it is shown for g-ratios in the range 0.5-0.75, the Rushton model performs best. Performance is only better for the outer diameter model for large g-ratios (between 0.75 and 0.95). Given that most axons conform to the former range of g-ratio, the Rushton model is the preferred approach, and thus an estimate of both inner diameter and g-ratio is valuable for mapping CV.

**Table 2:** Goodness of fit statistics for candidate simplified models to data generated from the Richardson model (*n* = 491, 520).

| Model | SSE | $R^2$ | Adj. $R^2$ | RMSE | $k$ | AIC | BIC |
| --- | --- | --- | --- | --- | --- | --- | --- |
| Rushton model | $1.54 \times 10^7$ | 0.766 | 0.766 | 5.69 | 1 | 1.76 | 12.87 |
| Linear outer diameter model | $1.58 \times 10^7$ | 0.760 | 0.760 | 5.77 | 1 | 1.76 | 12.87 |
| Polynomial expansion (full) | $1.23 \times 10^7$ | 0.813 | 0.813 | 5.09 | 11 | 21.81 | 144.0 |
| Polynomial expansion (cross-terms only) | $1.25 \times 10^7$ | 0.810 | 0.810 | 5.13 | 3 | 5.81 | 39.13 |

More complex models of CV derived using a polynomial expansion gave better fits than the two simpler models, but the parsimony measures suggest that, due to the increased number of coefficients, these models are suboptimal. Figure 4, shows the main improvement in the polynomial models is where AD is high and g-ratio is low. However, such axon configurations are uncommon: large diameter axons are unlikely to have very thick myelin sheaths. Therefore, there is little value gained by employing these more complex models to estimate CV *in vivo*.

In terms of sensitivity to errors in parameter estimation, we investigated the effects of bias in MRI-derived parameters, the sensitivity of CV estimates to these errors, and the sensitivity to noise. Overall we show that the errors in CV estimates are below 5% over a large region of the parameter space that applies most myelinated white-matter axons. CV estimates were least accurate when the AVF is small (below 0.3) and the AD is above 10 µm although less restrictive where AVF is high (greater than or equal to 0.5). This is to be expected as more sparse axon populations will generate less signal and reduce performance of model fitting. CV is most sensitive to errors in AVF, but errors in AVF are very small compared to other parameters. CV is most sensitive to errors in AD. This is as expected, since the Rushton model has highest sensitivity to AD.

While efforts have been made to incorporate true biological variability in the sensitivity analysis by taking parameter ranges from the literature, where available, the simulations are currently restricted to a single axon population. Variability should be considered across axon populations, where, for example, it is known that AD varies considerably throughout the CNS [46] as well as along single axons [47].

The computation of CV is assumed to be a valid aggregate measure of CV for a population of axons. The mean AD is parameterised by *λ* of the Poisson distribution and the g-ratio calculation has been shown to be valid for a distribution of ADs [23]. However, it is unclear if the CV value obtained from aggregated AD and g-ratio values is a valid aggregate representation of a distribution of CVs. The parametrisation of the AD distribution should be considered. A Poisson distribution was chosen as it has only one parameter, thereby reducing model complexity, but other distributions can offer better approximations of distributions observed in histology [48].

An assumption is made that the results of the axon simulations, whose baseline parameters are based on rat optic nerve [28], are generalisable to other white matter axons, and to other species. One aspect of this generalisation of particular concern is the g-ratio. There is a theoretical optimal g-ratio for a given fibre diameter [13]. In the present sensitivity analysis, the range of g-ratios tested is in an interval where the relationship between CV and g-ratio is monotonic and approximately linear. However, for other fibre populations with different ranges of g-ratios where the effect is non-monotonic, the sensitivities may differ substantially. Other parameters may present non-monotonic behaviour in parts of the parameter space not examined here. A potential future research avenue is to repeat the sensitivity analysis on a range of axon populations to see which populations better lend themselves to be modelled with MRI. However, obtaining all the morphological and electrophysiological parameters for multiple populations present significant practical challenges.

It has been demonstrated that the relative thickness of the water and lipid layers in myelin vary with age [39], and consequently the assumed constancy of *ω* used in Eq. 5 may not be valid. This issue can potentially be resolved by combining multiple myelin-sensitive contrasts, e.g. by adding in quantitative magnetization transfer (qMT). While qMT does not provide unique sensitivity to lipids, it does have sensitivity to protons bound to lipids and macromolecules. It may therefore be possible to exploit qMT and relaxometry methods together to better characterise the water-lipid ratio in myelin.

While this study explored the impact on MRI noise on CV estimates, there are a range of other sources of confounding variance, such as motion and distortions due to eddy currents, field inhomogeneities. Other sources of error could impact on CV estimates to differing degrees. Methodological issues around the estimation of axon diameters also should be considered. The apparent inter-axonal diffusion perpendicular to axon (which is used to estimate AD) is orders of magnitude smaller than the apparent extra-axonal diffusion [32, 33, 35]. This presents a challenge to estimating ADs, and require acquisitions at high *b* values to ensure a non-negligible contribution from the intra-axonal space [35]. On clinical MRI systems with gradients of up to 70 mT/m, this is problematic. On such systems, ADs below 6 µm will not be detectable. However, on a high gradient system (300 mT/m), where high *b* values are achievable, this can be reduced to 2-3 µm. [49, 50, 34]. Adapting the parameterised AD distribution with this limit can allow modelling of ADs below this limit [51]. We do stress that measuring axon diameter is challenging and we are not suggesting that it is possible to estimate CV everywhere within the brain [33]. An alternative to estimating internal AD is the framework of [52, 53, 54, 32, 33] allows characterisation of diffusion in the extra-axonal space in terms of the packing geometry of axons, which is dependent on the outer fibre diameter. This is appealing as this is closely correlated to CV, and only requires estimation of one microstructural parameter, instead of two, as used by the Rushton model. [33] suggest that the apparent correlation between AD and CV observed by [26] is due to contributions from the extra-axonal diffusion to the signal not being modelled correctly. However, it is unclear how outer fibre diameter can be disentangled from the packing geometry such as packing density and packing randomness within this framework. Also, as highlighted in the present study, the Rushton model is more accurate than the outer diameter model for estimating CV in more common ranges of AD and g-ratio. However the merits of this modelling framework should be explored further.

In conclusion, we demonstrated the feasibility of estimating CV for ensembles of axons from their diameter and g-ratio, estimated from *in vivo* microstructural MRI. These estimates can provide valuable insights into white-matter physiology, that would otherwise not be possible.

## 6 Acknowledgements

This work was supported by a Wellcome Trust Investigator Award (096646/Z/11/Z) and a Wellcome Trust Strategic Award (104943/Z/14/Z). The human MRI data were acquired at the UK National Facility for In Vivo MR Imaging of Human Tissue Microstructure funded by the EPSRC (grant EP/M029778/1), and The Wolfson Foundation.

We thank Lee Cossell & David Attwell for providing the electrophysiology simulation software used [28] and for guidance in its use. Thanks to Tobias Wood for valuable advice on the mcDESPOT processing, to Suryanarayana Umesh Rudrapatna for assistance in the human data acquisition, and to Silvia de Santis & Yaniv Assaf for their input into the AxCaliber method.

## A Validation of axon model

To ensure the implementation of the “Model C” axon model [27] produces results consistent with [28], simulations were carried out with the baseline condition and some parameters varied as described in this paper. All other model parameters were the same as used in the main simulations except that the stimulus current was fixed at 3 nA, as done in [28]. The baseline condition produced a CV of 2.95 ms^−1^, consistent with [28]. The results for other parameter variations are shown in Table A.1 which are also consistent with those reported in [28].

**Table A.1:**
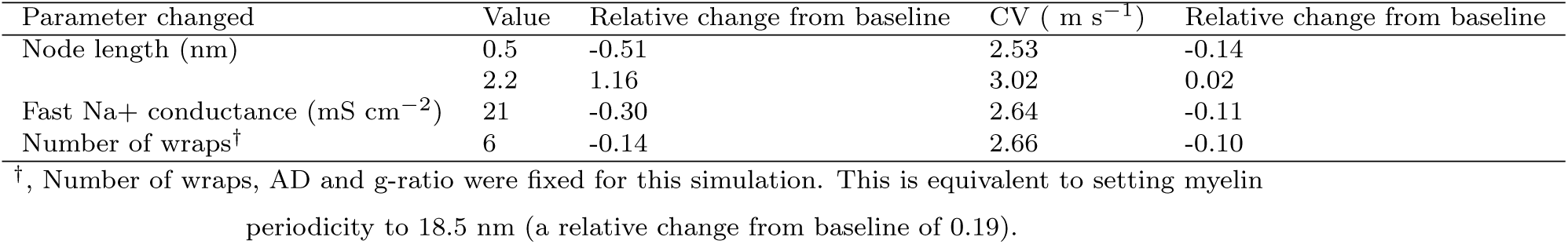
Changes in CV due to changes of parameters previously reported in [28].

| Parameter changed | Value | Relative change from baseline | CV ( $\text{m s}^{-1}$ ) | Relative change from baseline |
| --- | --- | --- | --- | --- |
| Node length (nm) | 0.5 | -0.51 | 2.53 | -0.14 |
|  | 2.2 | 1.16 | 3.02 | 0.02 |
| Fast Na<sup>+</sup> conductance ($\text{mS cm}^{-2}$) | 21 | -0.30 | 2.64 | -0.11 |
| Number of wraps<sup>†</sup> | 6 | -0.14 | 2.66 | -0.10 |

<sup>†</sup>, Number of wraps, AD and g-ratio were fixed for this simulation. This is equivalent to setting myelin periodicity to 18.5 nm (a relative change from baseline of 0.19).

## B OAAT sensitivity analyses

A one-at-a-time (OAAT) sensitivity analysis was performed for each parameter at 10 equally-spaced intervals within a ±20% range around the baseline condition. Results are shown in Figure B.1. It shows all sensitivity to all parameters is approximately linear over the interval tested.

In the main analysis, three of a set of six interdependent geometric parameters were manipulated: axon diameter, g-ratio and myelin periodicity. Three other parameters depend directly on these parameters: number of myelin wraps, myelin width and outer fibre diameter. As the impact of variance in these parameters will vary depending on which combinations of these parameters are fixed, we repeated the OAAT analyses for different combinations of fixings. The results are shown in Figure B.2. For most combinations, the sensitivity to each parameter shows a trend in the same direction regardless of which combination of other parameters are fixed. Notable exceptions are for the g-ratio where the direction of the sensitivity varies depending on which parameters are fixed. Sensitivity to g-ratio shows a negative trend when axon diameter and myelin periodicity or axon diameter and number of wraps are fixed. Sensitivity to g-ratio shows a positive non-linear trend when myelin periodicity and myelin width are fixed.

**Table C.1:** *In vivo* measurements of SNR and corresponding s.d. values for acquisition sequences used for estimating CV. S.D. values quoted for mcDESPOT scans are the effective noise added to recreate SNR measured in vivo.

| Sequence | Flip angle (°) | SNR | noise s.d. |
| --- | --- | --- | --- |
| CHARMED | 90 | 2,690 | 0.019 |
| AxCaliber | 90 | 1,776 | 0.024 |
| mcDESPOT - SPGR | 3 | 2,246 | 0.134 |
|  | 4.5 | 2,701 | 0.087 |
|  | 6 | 2,805 | 0.069 |
|  | 7.5 | 2,670 | 0.064 |
|  | 9 | 2,394 | 0.062 |
|  | 12 | 1,873 | 0.056 |
|  | 15 | 1,044 | 0.068 |
|  | 18 | 495 | 0.098 |
| mcDESPOT - SSFP | 10 | 10,136 | 0.010 |
|  | 15 | 14,256 | 0.008 |
|  | 20 | 17,666 | 0.008 |
|  | 25 | 19,090 | 0.007 |
|  | 30 | 19,659 | 0.007 |
|  | 40 | 18,269 | 0.007 |
|  | 50 | 14,272 | 0.008 |
|  | 60 | 9,776 | 0.010 |

## C SNR measurements

SNR for the diffusion sequences used for the AxCaliber and CHARMED acquisitions and the SPGR and SSFP acquisition used for mcDESPOT were measured by acquiring two image volumes: (1) a standard image volume (for diffusion sequences, only a *b* = 0 s mm^−2^ image was used as the noise distribution is not expected to be affected by the level of diffusion weighting) with the standard flip angle (see Table 3); and (2) a noise image volume acquired with exactly the same parameters but with the flip angle set to 0, such that there is effectively no echo received by the receiver coil. The noise s.d. was computed from the SNR. The results are shown in Table C.1

## D Estimating MVF from MWF

mcDESPOT gives a myelin volume fraction (MWF) that is the proportion of the total water volume that resides in the myelin layers.

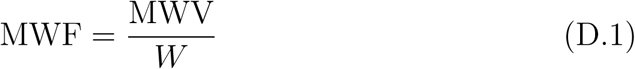

whereas calculation of g-ratios requires the myelin volume fraction (MVF), which is the proportion of the total volume that is myelin (both water and lipid components):

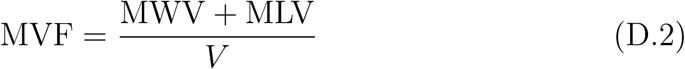

where MWV is the myelin water volume, MLV is the myelin lipid volume, *W* is the total volume of water and *V* is the total volume. If we assume that the ratio of water and lipid in the myelin compartment, *ω*, is constant, we can also express the MLV as:

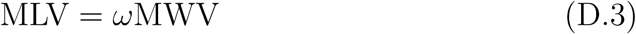

the total volume is given by:

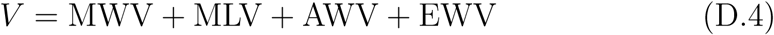

where AWV and EWV are the volumes of axonal and extracellular water, respectively. The total water is given by:

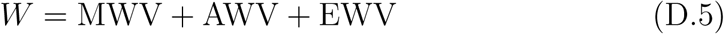

Assuming the only non-water compartment is the myelin phospho-lipid layers and that the contribution of phospho-lipid membranes from other cell types is negligible, this can be expressed as:

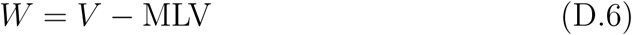

Substituting this into the expression for the MWF gives:

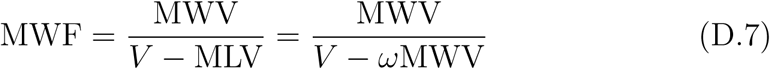

Collecting terms of MWV and rearranging gives:

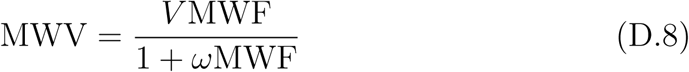

We can get an equivalent expression for the MLV by multiplying through by *ω*

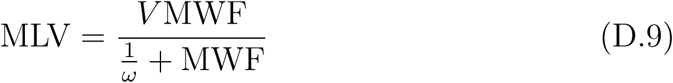

Substituting these into the expression for MVF gives:

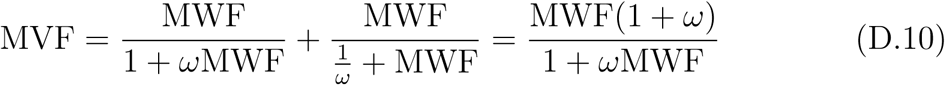

The value of *ω* was taken from [39] where the width of the lipid bilayers was about 4.6nm and the intra- and extracellular water layers were both 3.2nm, giving a lipid-water ratio of *ω* = 1.44. The relationship between MWF and MVF is show in Figure D.1.

**Figure B.1:**
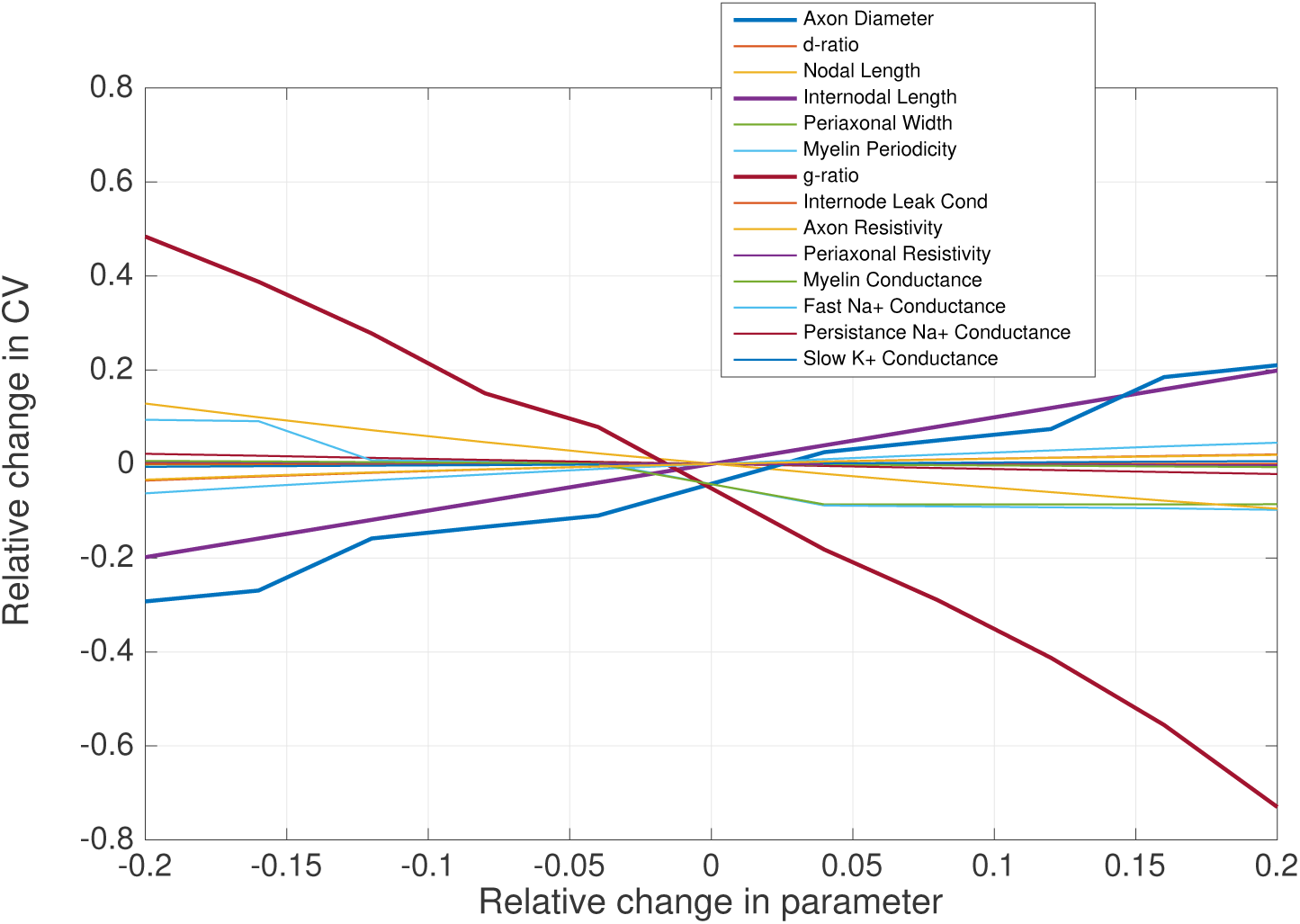
Results of OAAT analysis of sensitivity of CV to each of the free parameters tested.

**Figure B.2:**
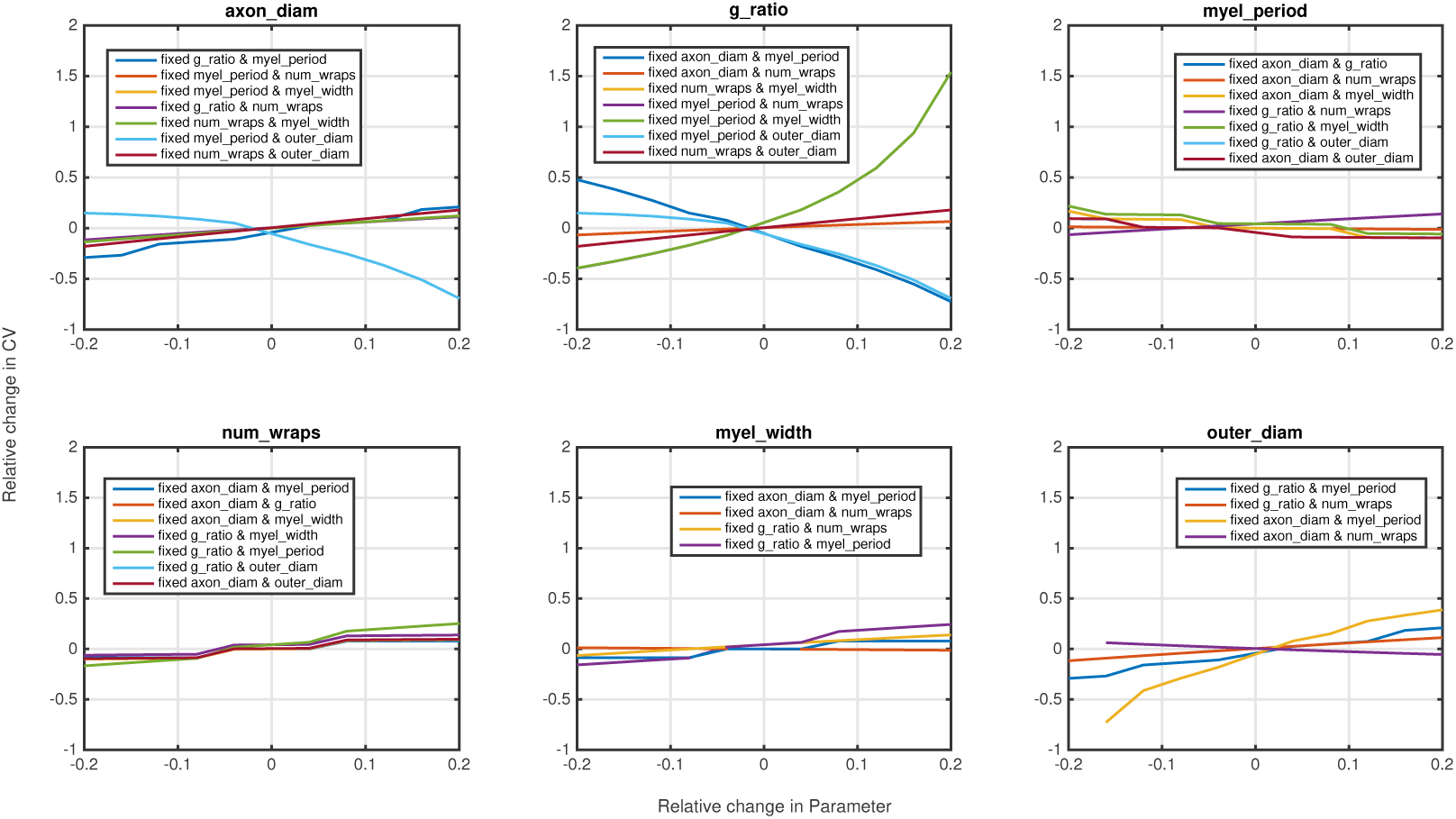
Results of OAAT analysis for the 6 interdependent geometric parameters with different combinations of parameter fixings.

**Figure D.1:**
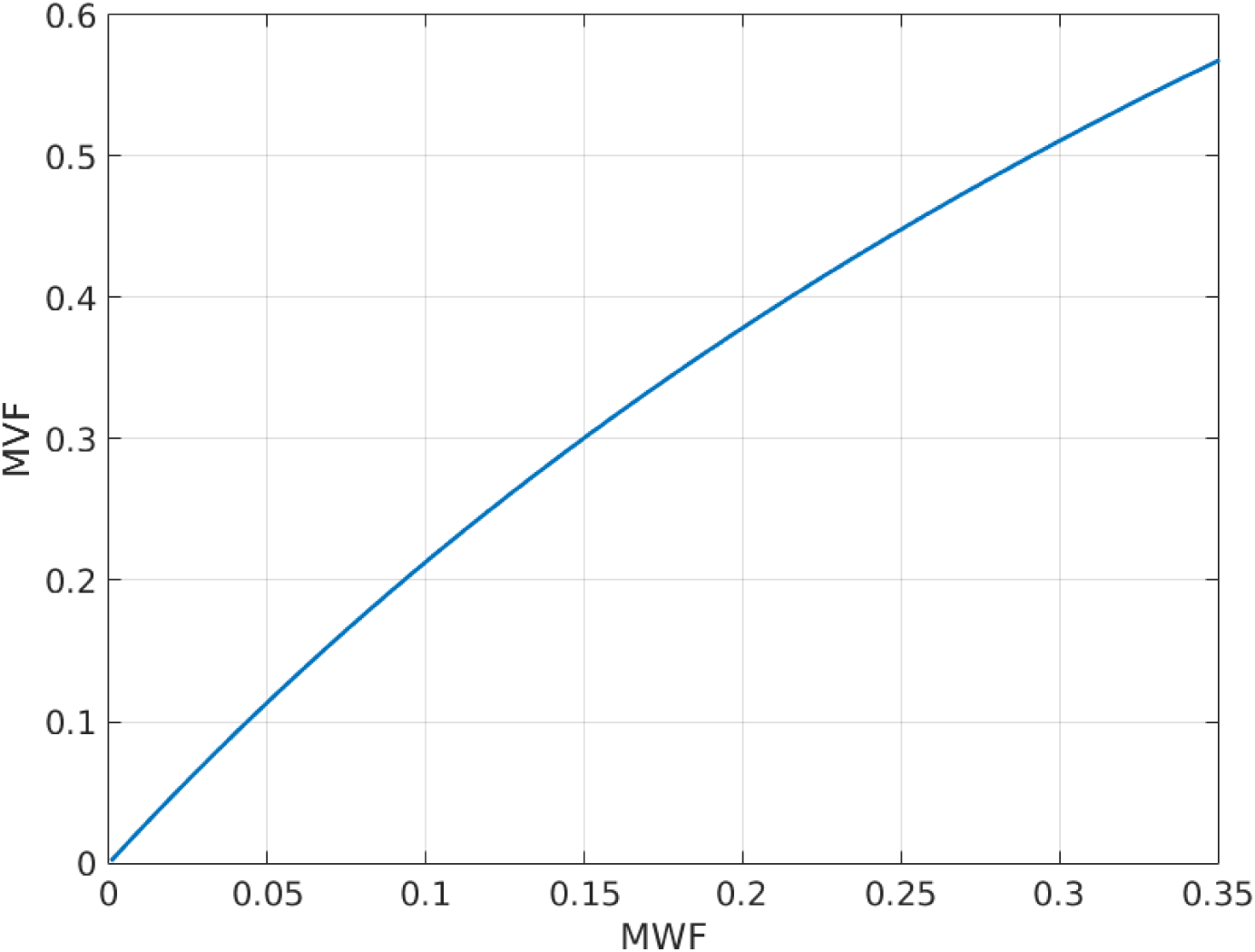
Theoretical relationship between MWF and MVF according to Eq.5.

